# Expanding the DNA Motif Lexicon of the Transcriptional Regulatory Code

**DOI:** 10.1101/2025.07.09.662874

**Authors:** Jingyu Fan, Virendra K. Chaudhri, Deepa Bisht, Nicholas A. Pease, Swapnil Keshari, Peter Gerges, David Reed Hall, Vivian Fang Yi, Susan Kales, Ching-Huang Ho, Jishnu Das, John P. Ray, Ryan Tewhey, Clifford A. Meyer, Nidhi Sahni, Harinder Singh

**Author notes:** co-corresponding authors: Clifford A. Meyer, Nidhi Sahni and Harinder Singh.

## Abstract

Cis-regulatory sequences in metazoan genomes contain elaborate combinations of transcription factor (TF) motifs. Stereospecific arrangements of simple motifs constitute composite elements (CEs) that enhance DNA-protein interaction specificity and enable combinatorial regulatory logic. Despite their importance, CEs remain underexplored. We integrate genome-scale computational discovery (CEseek) with customized massively parallel reporter assays (MPRAs) to uncover cell type-specific and shared CEs, corroborated by coincident and/or cooperative TF binding *in vivo* and *in vitro*. A deep learning model, GRACE, trained on MPRA data at single-nucleotide resolution, learns the CE lexicon and predicts activities of variants and alternative stereospecific CE configurations not included in the experimental design. Comparative analysis with models trained on orthogonal features such as chromatin accessibility, demonstrates convergence within the CE lexicon while resolving how individual motifs contribute to transcription versus chromatin accessibility. The expanded lexicon enables CE-aware predictions and interpretation of the impact of disease-associated non-coding variants in the human genome.

## Introduction

DNA-protein interactions mediated by transcription factors (TFs) decode regulatory information in metazoan genomes^1–5^. Structurally related TFs bind simple DNA motifs (6-10 bp) that have been extensively cataloged^3,6^ and constitute the primary lexicon of the gene regulatory code. Such catalogs are widely used to infer TF-to-target relationships and to build cell type- or cell state-specific gene regulatory networks (GRNs)^7–10^. However, metazoan promoters and enhancers consist of intricate combinations of simple motifs, generating a context-dependent regulatory syntax that remains challenging to decipher^2,5^ and that determines the spatiotemporal and signaling-responsive properties of regulatory DNA.

Transcriptional regulatory sequences contain composite elements (CEs) formed by precise juxtaposition of simple motifs that enable cooperative binding of TFs from distinct structural families. Exemplars include the ETS::IRF CE (EICE) recognized by PU.1/SPIB and IRF4/IRF8^11^; the AP-1::IRF CE (AICE) recognized by AP-1 (BATF/JUN) and IRF4/IRF8^12^; the NFAT::AP-1 CE bound by NFAT and AP-1 paralogs^13,14^; and the SOX::OCT CEs involving OCT4 and SOX family members^15^. Because CEs are stereospecifically constrained by the orientation and spacing of their constituent motifs, they carry greater information content and specificity in protein-DNA interactions and thus represent a higher-order lexicon of the gene regulatory code, with the potential to encode spatiotemporal transcriptional logic and enhance contextual specificity within GRNs. Despite their importance, the prevalence and diversity of CEs remain underexplored.

Previous studies of CE prevalence have relied on high-throughput *in vitro* DNA-binding assays with recombinant proteins (e.g., CAP-SELEX) or computational analyses of accessible or TF-bound chromatin regions^16,17^. Advances in chromatin accessibility profiling, massively parallel reporter assays (MPRAs)^18^, and interpretable deep learning^19–21^ now provide an opportunity to systematically discover, functionally validate, and computationally generalize CEs across cellular contexts. Herein we describe an integrated CEseek-MPRA-GRACE framework to achieve these goals.

## Results

### An integrated framework for discovery, testing and learning of CE lexicon

We hypothesized that CEs are diverse, widespread, and constitute a distinctive higher-order lexicon of the gene regulatory code and developed an integrated framework to test this (Fig. 1a). We designate enriched stereospecific motif configurations in cis-regulatory DNA as candidate CEs (cCEs) and reserve the term CE for those supported by experimental evidence, namely non-additive and/or configuration-dependent activity. CEseek performs genome-scale discovery of stereospecifically enriched motif pairs; the resulting cCEs are functionally tested by massively parallel reporter assays (MPRAs) with a custom TF-motif library; transcriptionally active CEs are evaluated against orthogonal evidence of TF co-binding *in vivo* and cooperative binding *in vitro*; and an interpretable deep learning model, GRACE, is trained on the MPRA data to predict activity at single-nucleotide resolution and infer the contributions of simple motifs and CEs. Comparison with orthogonal models trained on chromatin accessibility and histone modifications broadens evaluation of the encoded regulatory activities. To enhance accessibility, we developed the Transcriptional Composite Element Portal (CE portal; https://pitt-csi.shinyapps.io/ceportal/) for exploring the expanded lexicon.

**Fig. 1.**
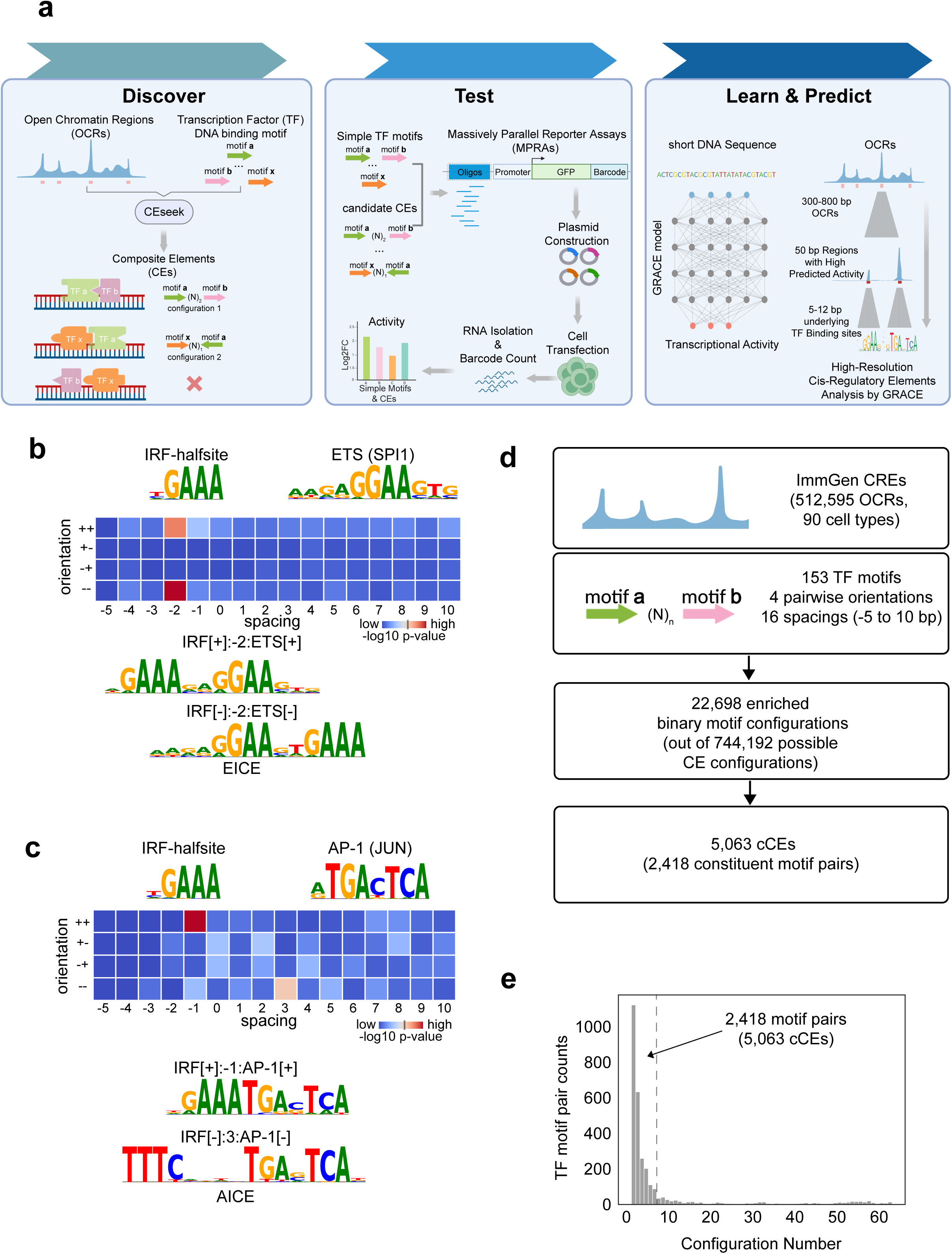
An integrated framework for discovery, functional testing, and deep learning of composite element grammar. **(a)** Schematic of the experimental and computational framework **Discover** - CEseek identifies candidate composite elements (cCEs) by searching for statistically enriched binary motif configurations by systematically varying the spacing and orientation of simple TF motifs in open chromatin regions (OCRs). Spacing between motifs is denoted as (N)_x_, indicating x base pairs. **Test** *-* Massively Parallel Reporter Assays (MPRAs) measure transcriptional activity of cCEs and the simple motifs comprising them along with their mutated counterparts. **Learn and Predict** - GRACE, a deep learning model trained on MPRA datasets, predicts transcriptional activity of CEs and simple motifs at single-nucleotide resolution. GRACE is integrated with orthogonal models (ChromBPNet, AlphaGenome) to evaluate CE contributions across chromatin accessibility, histone modifications, and diverse tissue contexts. **(b, c)** Heatmaps depicting the statistical enrichment of the indicated binary motif configurations (varying in orientation and spacing of the indicated simple motifs) in B-cell CREs. The enrichment was calculated using Fisher’s exact test with dinucleotide-shuffled sequences as the background, with a significance cutoff of −log_10_(p) > 20 indicated on the colorbar. The top panel **(b)** shows ETS::IRF (EICE) configurations, and the bottom panel **(c)** shows AP-1::IRF (AICE) configurations. Examples of the top two statistically enriched configurations of ETS::IRF and AP-1::IRF CEs that include well-characterized EICE and AICE composite motifs are displayed alongside the heatmaps. Sequence logos for IRF[-]:-2:ETS[-] configurations are presented as reverse complements for consistency in motif orientation. **(d)** Workflow for the discovery of cCEs in diverse immune and hematopoietic cell types as well as states. A total of 512,595 OCRs across 90 immune and hematopoietic cell types from the ImmGen ATAC-seq peak database were scanned for 153 non-redundant simple TF motifs using CEseek. All four pairwise orientations allowing for spacings of −5 to +10 bp were analyzed for occurrence in cell type/state-specific cCREs, resulting in 22,698 cell type-enriched cCEs (see **Methods**). Further filtering based on stereospecific constraints yielded 5,063 cCEs. **(e)** Histogram displaying the distribution of TF motif pair counts (n=2,418) with their numbers of statistically enriched configurations (n = 1 to 64). Significant motif pairs with ≤6 enriched configurations (n=5,063) out of a possible of 64 were deemed to be stereospecifically constrained (**d**) and selected as cCEs for further analysis.

### Analysis of candidate composite elements using CEseek

CEseek discovers cCEs as statistically enriched, stereospecific configurations of two simple TF motifs within cis-regulatory elements (Fig. 1a). It enumerates motif-pair configurations across user-defined spacing windows by scanning foreground open chromatin regions (OCRs) against matched background sequences, reporting enriched configurations with position weight matrices (PWMs) derived from aligned genomic instances. For non-palindromic motifs, each spacing yields four relative orientations, enabling comprehensive exploration of configuration space while retaining PWM interpretability.

We evaluated CEseek on murine B-cell enhancers previously characterized by STARR-seq^22^, scanning ETS, AP-1 and IRF motifs (IRF as a half-site) across a default spacing window (−5 to +10 bp) (Extended Data Fig. 1a). CEseek recovered the canonical ETS::IRF (EICE)^11^ and AP-1::IRF (AICE)^12^ composite motifs with the expected orientation and spacing constraints, including multiple stereospecific configurations consistent with known biochemical and functional properties (Fig. 1b,c). Thus, enrichment-based discovery recovers known composite motif syntax directly from endogenous regulatory DNA.

### Discovery of cCEs within immune and hematopoietic CREs

Next, we performed a genome-scale CEseek scan of the ImmGen murine cis-regulatory atlas^23^, a uniformly processed catalog of OCRs with exceptional depth spanning 90 well-defined immune and hematopoietic cell types and states (Fig. 1d). We used ImmGen data because it currently represents the most comprehensive and uniformly processed atlas of primary immune and hematopoietic chromatin accessibility, with no comparable human resource of similar breadth and cellular resolution presently available. We scanned with a curated collection of 153 representative simple TF motifs chosen to span major vertebrate TF DNA-binding specificities while minimizing redundancy; these covered ~78% of JASPAR 2024 Core Vertebrate motif clusters^24^ and captured the major immune and hematopoietic families (ETS, IRF, AP-1, NF-κB, RUNX, GATA, EBF, NFAT, KLF, C/EBP, PAX, E2A), although less common structural classes are underrepresented (Supplementary Note 1; Supplementary Table 1). Overall, we evaluated >10,000 motif pairs, each across 16 spacings (−5 to +10 bp) and 4 orientations (~750,000 configurations), allowing partial motif overlap because flanking base pairs carry lower information content^25^, cooperative binding can alter TF DNA-binding specificity^16^, and overlapping motifs can mediate anti-cooperative binding^26,27^.

Scanning cell-type- or state-specific OCRs (top 5%) against all other OCRs in the ImmGen atlas identified >20,000 enriched configurations (Fig. 1d; Supplementary Note 2). Consistent with our hypothesis, most motif pairs were enriched in only a small subset of possible configurations (Fig. 1e), yielding a bimodal distribution that separated pairs with conserved stereospecific configurations from those lacking such constraints (Extended Data Fig. 1b). Pairs with ≤6 enriched configurations were deemed “digital” and selected as cCEs for functional analysis (Supplementary Table 1); as expected, these were longer than their constituent simple motifs (Extended Data Fig. 1c,d) and included canonical CEs such as EICE^11^, AICE^12^ and NFAT::AP-1^13^, indicating that many digital cCEs represent new CEs within immune and hematopoietic CREs. CEseek also revealed a smaller set of “fuzzy” motif pairs with minimal spacing constraints and distinct mechanistic implications (see Discussion). cCEs are named based on the simple motifs used in the CEseek scan along with their orientations (+/-) and spacing/overlap (base pairs). Names of motifs do not specify the interacting TFs given that a motif can be recognized by multiple members within a structural family.

### Functional testing of cCEs using a custom MPRA library

To test cCE transcriptional activities, we designed a custom TF-motif MPRA library of ~4,000 cCE configurations derived from 2,418 motif pairs, each embedded in 30-bp endogenous sequences and assayed with deep barcode replication (~100 barcodes/oligo shared across all biological replicates; Supplementary Note 3; Supplementary Table 2). For 1,295 motif pairs, the library contained multiple stereospecific configurations (n=2-6), enabling direct testing of spacing and orientation effects on cCE activity. Given experimental constraints (Supplementary Note 3), the library was designed to sample biologically enriched CE configurations, rather than exhaustively test all theoretical configurations of each TF motif pair. Each cCE included four genomic instances with matched mutant controls disrupting one or both motifs (Fig. 2a), allowing quantification of motif activities and their non-additive functioning by comparing wild-type or single-mutant sequences with double-mutant controls in matched flanking contexts. The library also included single simple motifs and their mutants as qualitative references, together with 1,000 random 30-bp sequences as an independent activity baseline (Supplementary Note 4).

**Fig. 2.**
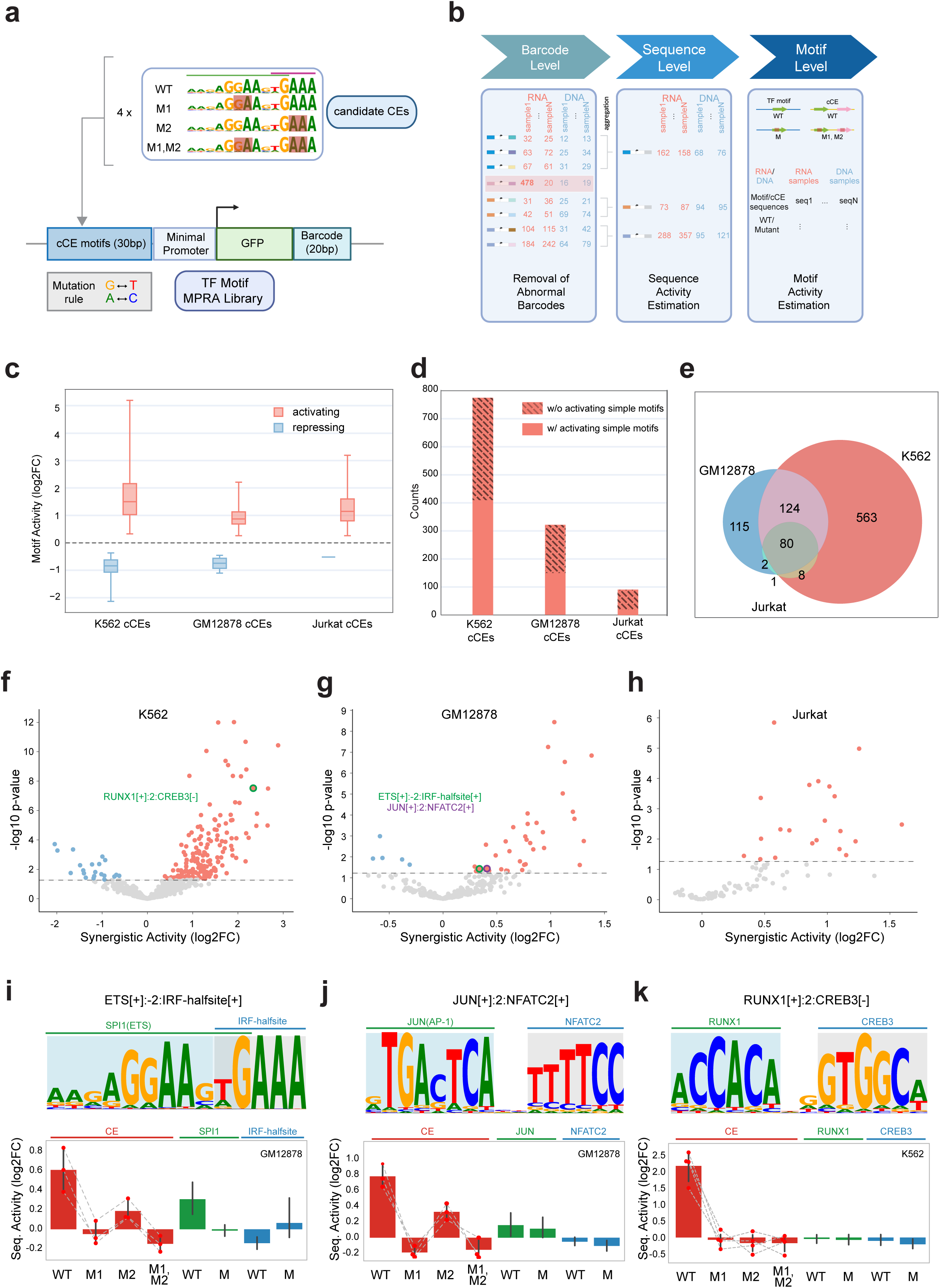
Functional testing of cCEs using a custom TF-motif MPRA library. **(a)** MPRA library design. The MPRA library comprises 30 bp oligonucleotides containing cCE configurations (n=3,795 configurations derived from 2,418 motif pairs, with 4 genomic instances of each). For 1,295 motif pairs, two to six configurations were included, enabling experimental testing of stereospecific effects. For simple motifs, each wildtype (WT) sequence is paired with a mutated version (M). For cCEs, each wildtype sequence is paired with three mutated variants: left motif mutated (M1), right motif mutated (M2), and both motifs mutated (M1, M2). Mutations were introduced at two consecutive positions with the highest information content in each TF binding site. Specifically, A was changed to C, T to G, and vice versa. The library additionally contained simple TF motif sequences as qualitative references (n=153, with 20 instances of each). The complexity of the library was n=60,714 sequences, and the activity of each sequence was assayable with >100 linked barcodes (see **Methods**). **(b)** MPRA analysis pipeline. The analysis of the TF-motif MPRA datasets is performed at three levels: barcode, sequence, and motif. CE transcriptional activities are quantified by comparing wild-type CE sequences to their double-mutant and single-mutant counterparts within the same endogenous flanking context, ensuring that flanking sequence differences are a controlled factor rather than a confounder (see **Methods**). At the barcode level, sample N denotes an individual DNA or RNA MPRA replicate. Barcode counts are first filtered to remove anomalous barcodes with excessively high read counts. At the sequence level, barcode counts corresponding to the same 30 bp oligonucleotide (seq N) are aggregated, and sequence activity is quantified by comparing RNA and DNA counts using DESeq2. At the motif level, simple motif activities are assessed by comparing the RNA/DNA log2 fold change (log2FC) of the wildtype motif-containing oligonucleotides with the log2FC of matched mutated control oligonucleotides. The activities of composite motifs are similarly estimated by comparing the log2FC values of the RNA/DNA ratios of sequences containing the wild type cCE with those of their double-mutant counterparts (see **Methods**). **(c)** Distributions of cCE transcriptional activities measured by MPRA in human cell lines. The MPRA library was transfected into K562 erythro-myeloid cells (6 replicates), GM12878 B cells (5 replicates), and Jurkat T cells (5 replicates) (see **Methods**). Box plots show the distributions of log2FC (wild-type vs. double-mutant) for activating (red) and repressing (blue) cCEs. **(d)** Counts of activating cCEs across the three cell lines (totaling ~900 across K562, GM12878, and Jurkat). Shaded bars highlight activating cCEs composed of simple motifs that individually lack detectable transcriptional activity in the indicated cellular context, accounting for ~half of all activating cCEs. Single-motif sequences included in the library served as qualitative references (see **Methods**). **(e)** Shared and cell type-specific activating cCEs. Pie chart illustrating the numbers of shared and cell-type specific activating cCEs among K562, GM12878, and Jurkat cells. **(f-h)** Synergistic and antagonistic interactions among constituent TF motifs within cCEs. Volcano plots highlighting 215 cCEs that exhibit significant non-additive behavior across the three cell lines: 190 with synergistic activation (AND-type logic) and 25 with attenuating or antagonistic effects (AND-NOT logic). Highlighted examples (colored points) correspond to the CEs shown in (**i–k**). **(i-k)** Examples of activating cCEs exhibiting AND-type synergistic logic. (i) EICE (ETS::IRF) in GM12878 B cells; (j) NFAT::AP-1 in GM12878 B cells; (k) RUNX::CREB in K562 hematopoietic cells. Sequence logos (PWMs) represent cCEs; bar plots show transcriptional activities of wild-type oligonucleotides (WT) and their single-(M1, M2) and double-mutant (M1,M2) variants, connected by dashed lines.

Given the high conservation of simple TF motifs between mouse and human, and assuming similar conservation for many CEs, we assayed the library in human cell lines widely used for regulatory genomics (ENCODE)^28^: K562 (hematopoietic), GM12878 (B lymphocyte), and Jurkat (T lymphocyte). MPRA data were analyzed at barcode, sequence and motif levels using matched mutant controls to define motif-level effects (Fig. 2b; Extended Data Fig. 2a-c; Supplementary Note 5; Supplementary Table 3-5), separating activity driven by motif grammar from flanking-context effects and estimating the contribution of each constituent motif within a cCE. Transcriptional activity was quantified by comparing RNA barcode counts with the corresponding DNA plasmid barcode counts, after which barcode counts were aggregated to the sequence level for downstream analyses.

Aggregated across the three cell types, we identified ~900 activating cCEs and a smaller set of repressing cCEs, with activating elements showing a broader dynamic range, consistent with the minimal promoter used (Fig. 2c,d; Extended Data Fig. 2d; “Functional cCEs” in CE portal; Supplementary Table 4). Notably, nearly half of the active cCE sequences comprised simple motifs that were individually inactive when assayed as qualitative references (Fig. 2d), indicating that their activity is an emergent property of the composite rather than of either motif alone. Most functional cCEs (66-72%) derived from motif pairs assayed in multiple configurations (n=2-6); permutation testing identified 125 (K562), 69 (GM12878), and 40 (Jurkat) motif pairs (n=183 unique pairs; some shared across cell types) with statistically significant configuration-dependent activity differences (p < 0.05), demonstrating that spacing and orientation shape activity beyond simple motif co-occurrence (Supplementary Note 6; Supplementary Table 4). For this subset of functional cCEs (n=183), configuration-dependent activity provides evidence supporting their classification as CEs.

Notably, ~75% of activating cCEs exhibited cell type-specific activity: GM12878-specific CEs were enriched for B-lineage motifs (e.g., EBF, E2A, PAX5, NF-κB) and K562-specific CEs for erythro-myeloid motifs (e.g., GATA, RUNX, SPI1, NFE2) (Fig. 2e; Extended Data Fig. 2e), indicating that composite grammar encodes context-dependent specificity and can drive emergent, cell type-specific transcriptional programs.

### CEs encode AND- as well as AND-NOT regulatory logic gates

To identify CEs with non-additive (emergent) function, we compared the activity of each wild-type CE with the summed activities (log-scale) of its constituent motifs, estimated from single-motif mutant sequences (Supplementary Note 5; Supplementary Table 5). This analysis defined 215 non-additively functioning CEs; 190 synergistically activating and 25 attenuating or antagonistic. CEs are hereafter designated as CEs based on these activity criteria (Fig. 2f-h). Canonical CEs such as EICE (Fig. 2i) and NFAT::AP-1 (Fig. 2j) were among the synergistic activators in the human B-cell context, exemplifying AND-type logic, while a novel RUNX::CREB CE in K562 cells (Fig. 2k) illustrates how composite grammar can generate emergent, cell type-specific activity in which CE output exceeds the summed activities of its single-motif-mutant variants.

Conversely, CEs consistent with AND-NOT logic paired an active motif with a weak or inactive partner to repress net activity, yielding significant negative synergy, including cases where the CE fell below the activation threshold yet the negative synergy was highly significant, consistent with transcriptional “quenching” (Supplementary Table 5). AND-NOT CEs trended toward more compact inter-motif spacing than AND CEs in K562 and GM12878, consistent with steric exclusion, though this did not reach significance (Supplementary Note 7). Rarely, we observed XOR-like behavior in which pairing two active motifs attenuated net activity, indicating mutual interference (Supplementary Table 5). Together, composite motif pairings encode diverse two-input regulatory logic spanning cooperative activation, antagonism and repression.

### TF-focused analyses of CEs using CAP-SELEX and ChIP-seq datasets

We next evaluated functional cCEs for cooperative TF binding *in vitro* by matching their PWMs to those in CAP-SELEX^16,29^ (Supplementary Note 8; Supplementary Table 6; “CEs x CAP-SELEX CEs” in the CE portal). CEs with strong matches to their CAP-SELEX counterparts (Fig. 3a-c) provided supporting evidence for cooperative binding of specific TF pairs *in vitro*.

**Fig. 3.**
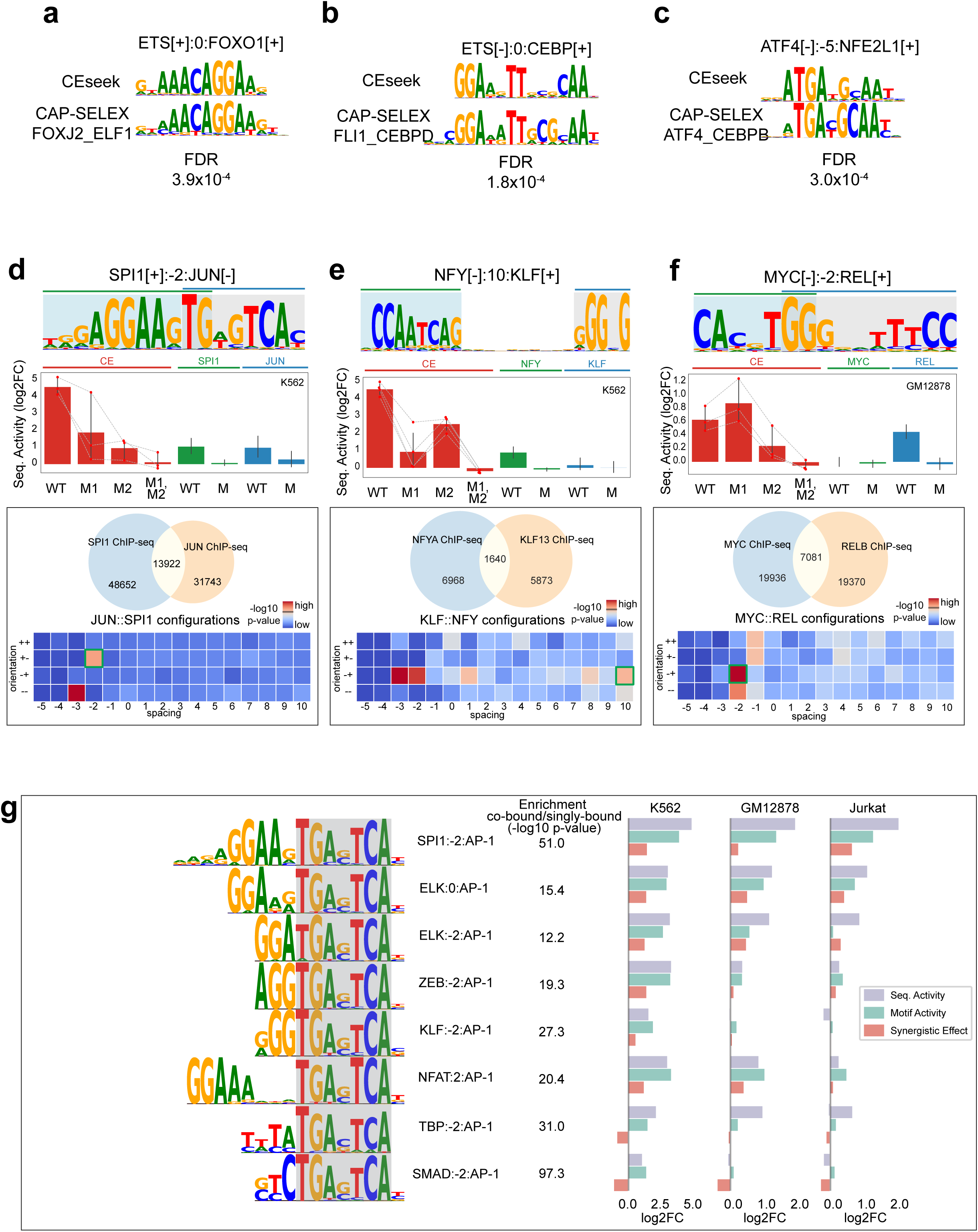
Analysis of CEs for cooperative TF binding in vitro (CAP-SELEX) and co-binding in vivo (ChIP-seq) **(a-c)** Representative CEs demonstrating significant matching with CAP-SELEX-derived cooperative binding motifs. For each CE, the sequence logo (PWM from CEseek) is shown alongside the matched CAP-SELEX PWM with its TomTom FDR value. **(d-f)** Representative CEs demonstrating enrichment in co-bound genomic regions of cognate TFs. The top panels display sequence logos of the CEs (PWMs), whereas the bar plots show the activities of the wild-type oligonucleotides and their mutated variants (connected by dashed lines) in the indicated cells. For each CE, cognate TF ChIP-seq datasets were identified from CistromeDB by TF family matching. Venn diagrams show co-bound versus singly bound peak regions for the relevant TF pair. CEseek was applied to co-bound regions using singly bound regions as background to assess configuration-specific enrichment; heatmaps depict the resulting enrichment of all 64 CE configurations (spacings × orientations), with the MPRA-validated configuration, corresponding to the PWM above, highlighted (green box). In panel (**d**), the top two enriched configurations are effectively equivalent due to the palindromic nature of the JUN motif. Panel (**f**) shows an example of an antagonistic CE (MYC::REL) with (AND-NOT) transcriptional logic. **(g)** Diversity of CEs comprising AP-1 motifs. Sequence logos of the AP-1 families of CEs are shown on the left. The bar plots show the measured sequence, motif, and synergistic activities of each CE in K562, GM12878, and Jurkat cells. Supporting evidence for TF co-binding was obtained from available CistromeDB ChIP-seq datasets for the specific TF family members listed in Supplementary Table 8.

To test whether MPRA-delineated cCEs are associated with co-binding of cognate TF pairs *in vivo*, we applied a three-step approach using CistromeDB^30–32^ ChIP-seq datasets (Supplementary Note 8; Supplementary Tables 7,8). We first matched TF motifs within CEs to ChIP-seq datasets for TFs of the same family; next assessed whether synergistic (activating or repressing) CEs were enriched within cognate TF ChIP-seq peaks; and finally tested, by CEseek, their enrichment specifically within co-bound regions using singly bound regions as background. Exemplar CEs enriched in co-bound regions are shown (Fig. 3d-g; Extended Data Fig. 3c), along with their configuration-specific enrichment. In many cases the MPRA-validated configuration ranked among the most enriched of the 64 possible configurations for that motif pair, supporting the stereospecific nature of composite binding *in vivo*. We also found antagonistic CEs enriched in co-bound regions, exemplified by MYC::REL (Fig. 3f), consistent with anti-cooperative binding or co-recruitment manifesting as transcriptional “quenching” (Supplementary Tables 6,9; “CEs x CistromeDB CEs” in the CE portal). Thus, a substantial fraction of MPRA-active CEs corresponded to composite binding sites promoting cognate TF co-occupancy *in vivo* and/or cooperative TF binding *in vitro*.

### Combinatorial diversity of CEs and their signaling logic

We next examined combinatorial diversity of CEs by identifying the most frequent simple motifs. AP-1, KLF and ETS motifs were most common (Extended Data Fig. 3a) and, by combining with distinct partners, greatly diversified the CE repertoire (Extended Data Fig. 3b). Representative AP-1 and KLF CEs are shown with their enrichment in regions co-bound by cognate TF family members (Fig. 3g; Extended Data Fig. 3c; Supplementary Note 9). AP-1 dimers of FOS, JUN, ATF and MAF are activated by growth factors, hormones and cytokines to control proliferation, differentiation and stress responses^33^, and their CEs predict integration of MAPK regulated AP-1 activity with distinct signaling pathways: SMAD::AP-1 with TGF-β, NFAT::AP-1 with calcium and ZEB::AP-1 with Wnt. These illustrate how the CE lexicon can encode combinatorial logic for integrating multiple signaling inputs into coordinated transcriptional responses.

### Learning CE grammar with GRACE

We developed GRACE (Genome Regulatory Analysis of Simple and Composite transcriptional Elements), an interpretable deep learning model trained on TF-motif MPRA measurements to predict transcriptional output from short sequences and resolve motif contributions at single-nucleotide resolution (Fig. 4a; see Methods). GRACE combines convolutional filters (learning motif-like patterns) with attention-based modeling of motif-motif interactions (transformer encoder) to represent stereospecific composite grammar; randomized flanking sequences mitigate the central-position bias of fixed 30-bp oligos, and locally restricted attention preserves single-nucleotide interpretability while supporting variable-length inputs. GRACE predictions correlated strongly with measured activities on held-out sequences, approaching replicate-level consistency (Fig. 4b; Extended Data Fig. 4a-e; Supplementary Note 10), and outperformed regression models, gapped k-mer SVMs (LS-GKM)^34^, and conventional CNNs (Supplementary Note 11).

**Fig. 4.**
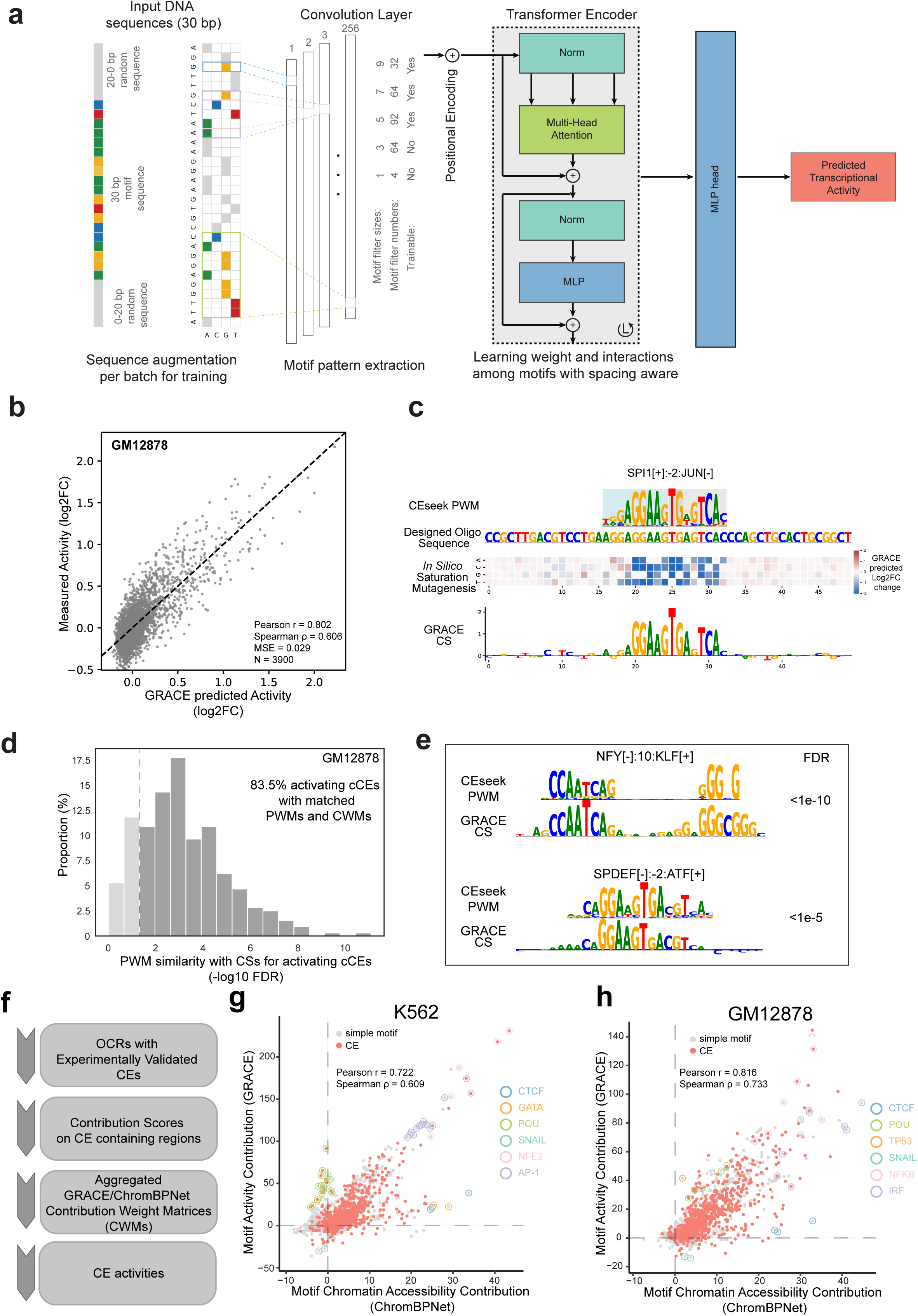
Learning CE grammar with GRACE. **(a)** Model overview. GRACE is a neural network model that integrates convolutional layers to learn TF binding motifs and employs self-attention with positional encoding to capture interactions among these motifs. See **Methods** for details. **(b)** Prediction accuracy of GRACE. Scatterplot comparing GRACE-predicted log2FC transcriptional activities for withheld test sequences (n=3,900) with their experimentally measured MPRA activities in GM12878 cells. The Pearson and Spearman correlations are indicated in the panel. (Training set: n=49,042 sequences; see **Methods**.) **(c)** Generating transcriptional activity contribution scores with GRACE. Illustration of GRACE-assigned nucleotide contribution scores via in silico saturation mutagenesis (ISM), shown for the SPI1[+]:-2:JUN[-] CE. The 30 bp oligonucleotide corresponding to this CE was systematically mutated; the ISM heatmap shows predicted log2FC for each single-nucleotide substitution. The resulting contribution score (CS; bottom) is compared with the CEseek PWM (top) to validate that GRACE recovers the composite motif at single-nucleotide resolution. **(d)** Histogram showing the TomTom FDR distribution for similarity between CEseek PWMs and GRACE-derived CWMs for activating cCEs, using the GM12878 GRACE model. The fraction of matching cCEs (FDR < 0.05) is indicated. **(e)** Examples of matched CEseek PWMs with GRACE predicted CWMs. The FDR values for each pair are indicated. **(f)** Workflow for CE motif contribution analysis. OCRs containing experimentally validated CEs are first identified. Model-derived contribution scores (from GRACE or ChromBPNet) are computed across CE-containing sequences. These scores are aggregated across motif instances to generate CWMs. Finally, CE activity is quantified based on normalized motif-level contribution scores derived from the CWMs. **(g, h)** Analysis of TF motif contributions to chromatin accessibility and transcriptional activity using ChromBPNet and GRACE models. Scatter plots comparing motif contributions to chromatin accessibility (x-axis; ChromBPNet) and transcriptional activity (y-axis; GRACE) for simple (gray) and CE (red) motifs in K562 (**g**) and GM12878 (**h**) cells. Contribution scores are normalized as fold-changes relative to 50 bp flanking region scores (see (**f**) and **Methods**). Representative motifs are annotated, including CEs with high transcriptional activity, CTCF (high accessibility, low transcriptional contribution), GATA (high accessibility in K562 but not GM12878), and POU-domain factors (high transcriptional contribution in both cell types).

Despite the library’s complexity (~4,000 cCEs with mutant and alternate-configuration variants), the experimentally accessible sequence space remained limited: the single-nucleotide variant space of a 30-mer and the 64 configurations per motif pair are prohibitively large (Supplementary Note 12). GRACE was therefore designed to learn from the sampled sequence space and generate testable predictions across the wider theoretical space.

As an initial application of GRACE, we performed *in silico* saturation mutagenesis (ISM) to identify the motifs underlying CE transcriptional activity (Supplementary Note 13), reasoning that if GRACE learns the CE lexicon at single-nucleotide resolution, its contribution weight matrices (CWMs) should resemble the CEseek PWMs, as shown for the SPI1[+]:-2:JUN[-] CE (Fig. 4c). Consistent with this, a large proportion of active CEs showed strong CWM-PWM similarity (Fig. 4d,e; Extended Data Fig. 4f,g; “CE ISM plots” in CE portal; Supplementary Table 10). Although trained on only ~50,000 sequences of the vast theoretical space (4³⁰) of a 30-mer, the designed TF-motif library enabled GRACE to capture key motif grammar across the sampled space, indicating that the transcriptional activities of many CE-containing sequences are dictated by the CEseek-identified motifs rather than unrelated features.

We next projected the GRACE-learned CE lexicon onto endogenous chromatin and transcription features in GM12878 B cells. GRACE-predicted activity across OCRs was strongly concordant with markers of active regulatory regions, including H3K27ac (Supplementary Note 14), and was spatially positioned within bidirectional transcription regions spanning promoters and enhancers mapped by GRO-cap, with inter-segment spacing at enhancers matching that of bidirectional initiation sites (Supplementary Table 11). Most MPRA-defined CEs fell within GRACE-predicted active enhancer segments, indicating that the CE lexicon corresponds to functional elements in their endogenous contexts.

To assess how the GRACE-learned CE lexicon contributes to chromatin accessibility, we used the orthogonal model ChromBPNet^21^, trained on ATAC-seq (Supplementary Note 15). In both models, motif contributions were quantified as the fold difference between aggregated base-resolution contribution scores on motif hits versus flanking regions (±50 bp) within OCRs (Fig. 4f; see Methods), evaluated for K562 and GM12878 OCRs using JASPAR 2024 CORE vertebrate motifs^24^ or the validated CEs. Many CEs showed strong correlations between predicted transcriptional activity and accessibility contributions (Fig. 4g,h; Supplementary Table 12), with consistent CWM patterns across models (Extended Data Fig. 4h,i; “CE contribution plots” in CE portal). Thus, convergence of CEseek-identified CE motifs with their GRACE transcriptional contributions and ChromBPNet accessibility contributions strongly supports the validity and biological relevance of the expanded CE lexicon. Nevertheless, direct benchmarking on held-out MPRA sequences showed that GRACE substantially outperformed ChromBPNet in predicting transcriptional activity (GM12878: Pearson r = 0.80 vs 0.60; K562: r = 0.81 vs 0.60; Extended Data Fig. 4k; Supplementary Note 16), consistent with the different regulatory features learned by the two models.

We next examined how simple and composite motifs differentially contribute to these modalities. CEs consistently ranked among the strongest contributors to both transcription and accessibility, whereas their constituent simple motifs contributed more modestly, the composite context amplifying predicted effects, likely through cooperative TF interactions. Informative differences between the models were also apparent (Fig. 4g,h). CTCF, though absent from the MPRA library, was learned by GRACE from related sequences and contributed modestly to transcription (Extended Data Fig. 4j) but was assigned much larger accessibility contributions by ChromBPNet, consistent with its architectural rather than transcriptional role^35^. Conversely, POU-domain factors showed strong transcriptional but limited accessibility contributions in both K562 and GM12878, suggesting action within already accessible chromatin. Cell-type differences also emerged: GATA motifs contributed strongly to accessibility in K562 but minimally in GM12878, consistent with lineage-restricted pioneer activity^36^.

We next used GRACE to predict activities for all 64 configurations of each motif pair constituting validated CEs in K562 and GM12878 (Supplementary Note 17; Supplementary Table 13). For configurations absent from the MPRA training set, GRACE predictions remained highly concordant with ChromBPNet (K562: Pearson R = 0.693, Spearman R = 0.601; GM12878: Pearson R = 0.633, Spearman R = 0.757), approaching the agreement seen for experimentally tested configurations (K562: R = 0.719/0.627; GM12878: R = 0.799/0.781) (Fig. 5a,b; Extended Data Fig. 5a-d; “CE configuration plots” in CE portal). Substantial convergence (Pearson R > 0.6) was observed for 64% of pairs in K562 and 58% in GM12878, defining a set of high-confidence CE configurations, exemplified by JUN::MAF (Fig. 5c,d). That this agreement extends to configurations not in the MPRA design indicates the short CE-containing sequences and their mutant variants provided sufficient implicit coverage for GRACE to learn the underlying composite grammar.

**Fig 5.**
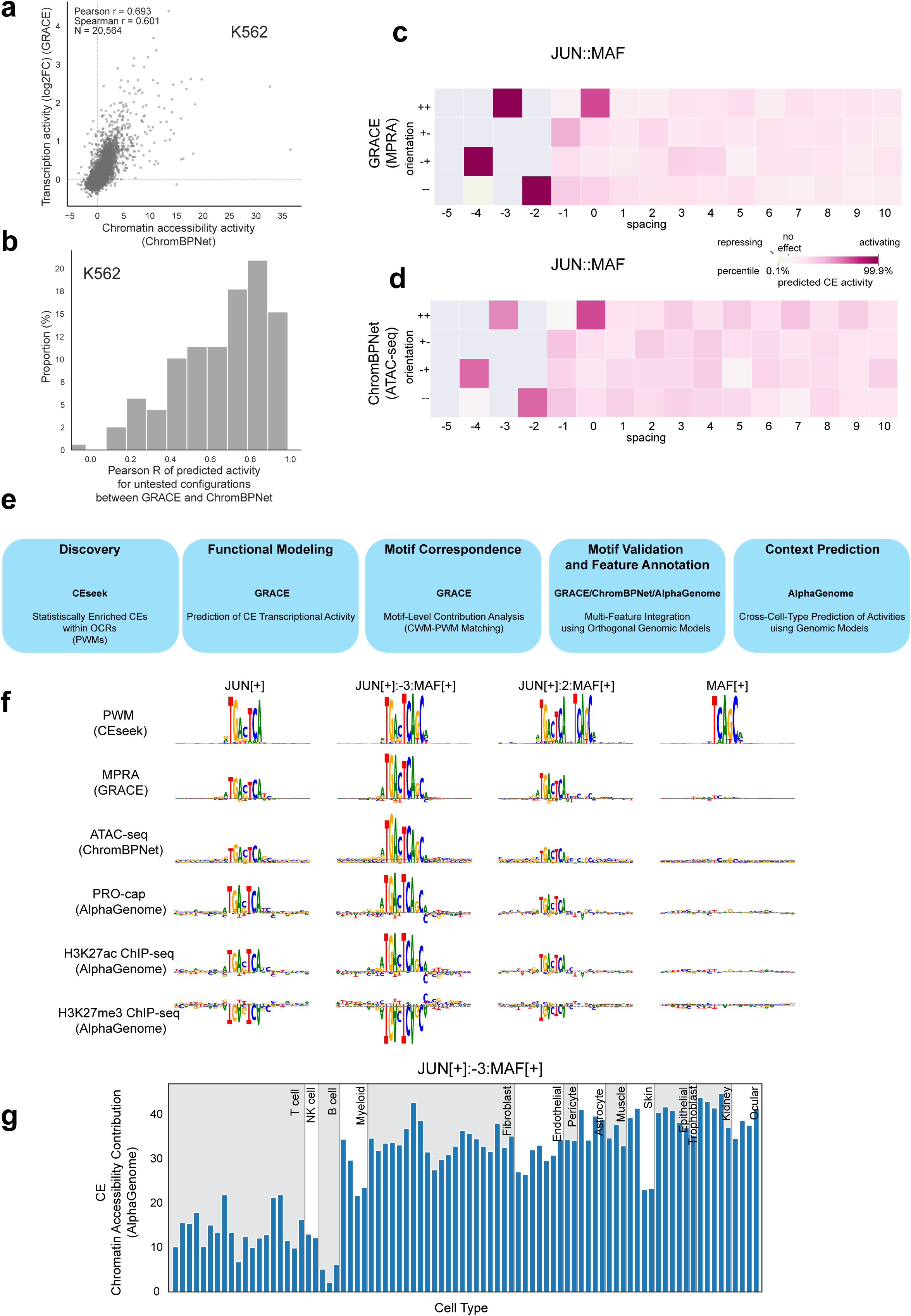
Generalization of GRACE to untested CE configurations and multi-modal regulatory analysis. **(a)** GRACE and ChromBPNet predictions for untested CE configurations. Scatter plot comparing GRACE-predicted transcriptional activity and ChromBPNet-predicted chromatin acces sibility for CE configurations not included in the TF-motif MPRA training set (K562 cells). GRACE configurations used in training were masked. **(b)** Histogram of Pearson correlation coefficients between GRACE and ChromBPNet predictions across individual motif pairs for untested configurations (K562 cells, median Pearson R = 0.710). **(c, d)** Activity predictions of stereospecific JUN::MAF CE configurations. Heatmaps show GRACE-predicted transcriptional activity (**c**) and ChromBPNet-predicted chromatin accessibility contributions (**d**) across all 64 configurations (4 orientations × 16 spacings). The JUN[+]:-3:MAF[+] configuration (functional CE validated by MPRA; see (**f**)) is highlighted. Color scales represent percentile of predicted activity within each model’s predictions respectively. **(e)** Schematic overview of the integrated framework for discovery, functional modeling, and validation of CE configurations. The workflow spans (i) CE identification using CEseek, (ii) transcriptional activity prediction with GRACE, (iii) motif-level correspondence through single-nucleotide GRACE contribution analysis, (iv) motif validation and multi-feature integration using orthogonal models and (v) cross-cell-type activity prediction. **(f)** Example of JUN::MAF CEs and their constituent simple motifs, illustrating motif contributions across multiple regulatory modalities. Shown are position weight matrices (PWMs; CEseek) and model-derived contribution profiles (CWM) from GRACE (MPRA), ChromBPNet (ATAC-seq), and AlphaGenome (PRO-cap, H3K27ac, H3K27me3), highlighting configuration-specific effects on transcriptional activity and chromatin features. The JUN[+]:-3:MAF[+] configuration (second column) represents a novel functional CE predicted by GRACE and supported by MPRA. The JUN[+]:2:MAF[+] configuration (third column) is a non-functional configuration included to illustrate stereospecificity: it shares the same motif pair but differs in spacing, producing no detectable activity contribution (see heatmaps in (**c**)). **(g)** Predicted chromatin accessibility contributions of the JUN[+]:-3:MAF[+] CE across diverse primary cell types and tissues, as inferred from AlphaGenome DNase-seq models. Each row represents a cell type or tissue in indicated groups; bars display predicted accessibility contributions.

### Extending CE analysis across tissues and regulatory modalities

We extended the analysis of CEs to incorporate additional molecular features and cellular contexts using AlphaGenome^37^, which integrates regulatory-genomics models trained on diverse transcription and chromatin features (Fig. 5e; Supplementary Note 18). AlphaGenome enabled assessment of CE contributions to multiple histone modifications, including H3K27ac and H3K27me3, revealing that CE configurations differ in their capacity to promote active versus repressed chromatin states, and corroborated CE function while resolving stereospecific contributions across modalities (Fig. 5f; Extended Data Fig. 5e). In K562, NFY::KLF and JUN::MAF pairs showed configuration-specific effects across alternative spacings, and the NFY::KLF CE made no discernible contribution to H3K27ac, in contrast to JUN::MAF. Leveraging AlphaGenome’s DNase I models across diverse primary tissues, we further inferred cell-type-specific activity landscapes of validated and predicted CEs beyond the hematopoietic and immune contexts assayed by MPRA (Fig. 5g; Extended Data Fig. 5f), extending the lexicon’s utility and providing a broadly applicable framework for CE prioritization in tissue-specific studies. Thus, CEs are prominent, interpretable contributors to both chromatin state and transcriptional output, and integrating models across modalities enables mechanistic dissection of CE function at endogenous loci.

### Using CE-aware modeling to interpret functional non-coding variants

We next applied GRACE to predict allele-specific effects of disease-associated non-coding variants, distinguishing those disrupting simple motifs from those affecting CEs. Using an MPRA of reference and alternate autoimmune-associated alleles in a human B-cell context^38^, GRACE predictions on variant-centered sequences were strongly concordant with their measured allele-specific effects, and precision-recall analyses showed high precision among high-scoring predictions (Fig. 6a,b; Supplementary Note 19; Supplementary Table 14), highlighting the utility of CE-aware modeling for interpreting noncoding variation.

**Fig. 6.**
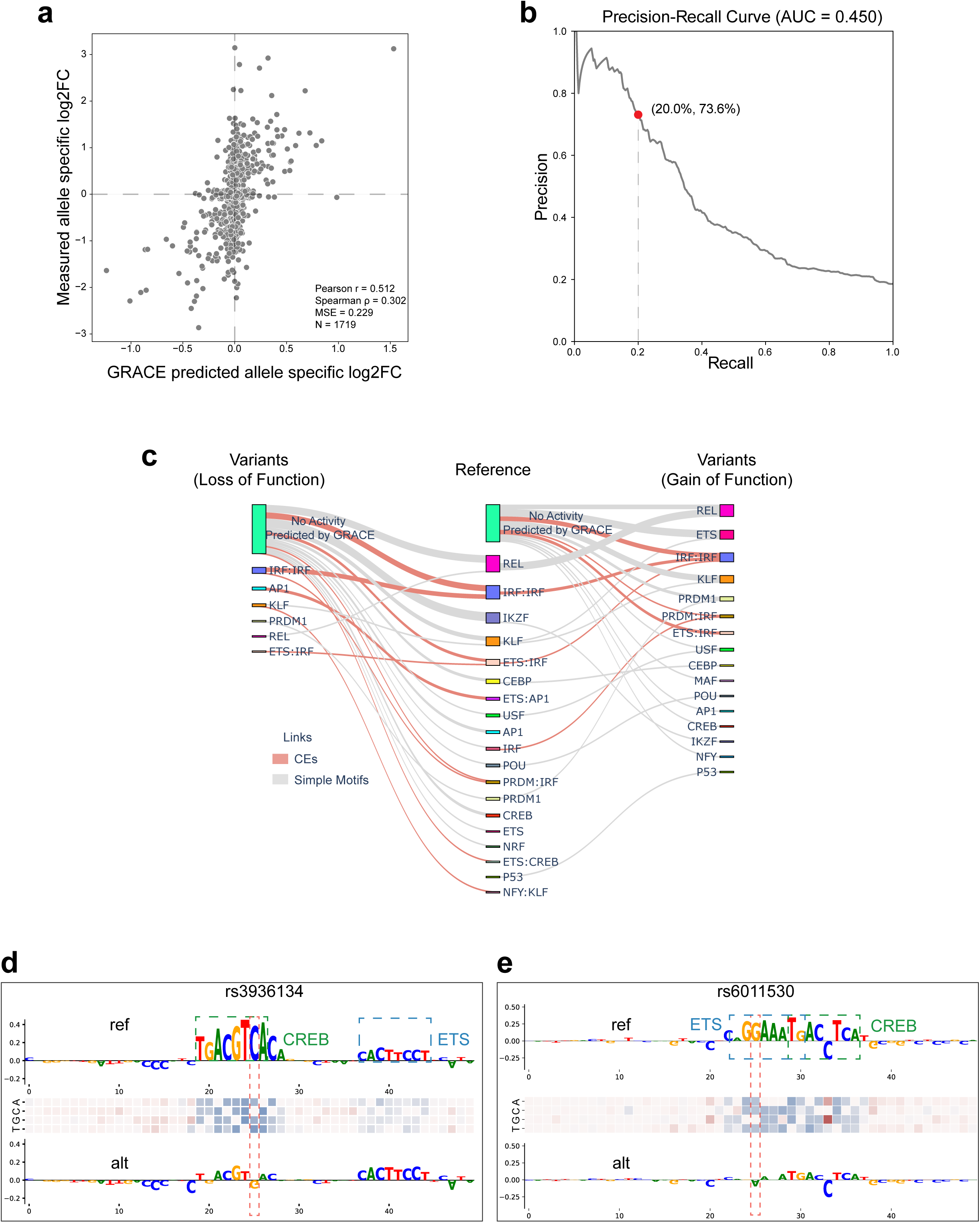
CE-aware GRACE model predicts allele-specific effects of autoimmune disease-associated GWAS variants. **(a)** GRACE predictions of GWAS variants. Scatter plot comparing GRACE-predicted and MPRA-measured allele-specific effects (log2FC) for 1,719 autoimmune GWAS variants (emVARs) assayed in an EBV-transformed B-cell line from a lupus patient (see **Supplementary Table 14**). **(b)** Precision-recall curve testing the performance of the GRACE model in identifying GWAS variants with significant allele-specific effects in the MPRA dataset. **(c)** Sankey diagram linking GRACE-interpreted motif features to predicted allele-specific transcriptional effects (gain- or loss-of-function, defined as |log2FC| ≥ 0.25) for n=68 variants from **Fig. 6a** for which GRACE assigned motif-level interpretations. CE motifs (red lines) and simple TF motifs (gray lines) are linked to their associated alleles. **(d, e)** Examples of autoimmune disease-associated GWAS variants. Sequence logos showing nucleotide contributions for the reference (top row) and alternative alleles (bottom row) with their annotated TF motifs. The red dashed line indicates the variant position and nucleotide substitution. The heatmap in the middle row of each panel displays *in silico* saturation mutagenesis predictions generated by GRACE for the reference sequences. Both alternative alleles are predicted to result in loss-of-function. The variant in (**e**) disrupts an ETS::CREB CE (rs6011530; SLE-associated); CEseek analysis supporting this CE configuration using CREB1 and SPI1 ChIP-seq co-binding in GM12878 cells is shown in **Extended Data Fig. 6a**.

Mechanistically, among variants that GRACE predicted and localized to specific motif features, a substantial fraction (30%) mapped to CEs (Fig. 6c-e; Extended Data Fig. 6a-e). In several cases, disruption of one constituent motif produced collateral effects on its partner within the composite (Fig. 6e), consistent with the CE behaving as a single functional unit; for example, one ETS::CREB CE configuration which was enriched in regions co-bound by SPI1 (PU.1) and CREB1 (Extended Data Fig. 6a). Gain-of-function variants also created *de novo* composite configurations (Extended Data Fig. 6d,e). We extended GRACE predictions to 5,118 autoimmune-associated variants (GM12878 models; “Variant impact contribution plots” in CE portal) curated from PICS2^39^ and the Pan-UK Biobank^40^, spanning >30 diseases including SLE, MS and RA (Supplementary Table 14), providing a community resource for prioritizing functional noncoding variation. Together, a substantial fraction of functional noncoding variation appears to operate through perturbation of composite grammar rather than isolated TF motifs and pairing the CE lexicon with CE-aware models that separately quantify transcriptional and chromatin effects provides a mechanistically informative framework for variant interpretation beyond single-motif annotations.

## Discussion

Metazoan enhancers and promoters contain dense combinations of simple TF motifs, yet most genome-wide annotations and GRN models treat motifs as independent features. Our results establish that composite elements constitute an extensive higher-order lexicon that is prevalent in regulatory DNA and functionally consequential. By integrating CEseek-based discovery, direct functional testing with a custom TF-motif MPRA, an interpretable deep learning model trained on the resulting data (GRACE), and orthogonal sequence-to-function models, we move from a “parts list” of simple motifs toward a grammar-aware transcriptional regulatory code.

CEseek delineated a digital subset of motif pairings with restricted orientations and spacings, consistent with structurally constrained TF-TF interactions. MPRA measurements showed that many digital configurations have strong transcriptional consequences, activating or repressing, and that the activity of a simple motif can vary depending on its composite partner and configuration. Across three cell types, we identified ~900 functional cCEs, among which 215 exhibited non-additive activity and 183 showed configuration-specific activity. These 339 CEs, satisfying one or both MPRA-based activity criteria (215 non-additive, 183 configuration-dependent, 59 satisfying both), substantially expand the functionally characterized regulatory grammar.

The composite motif lexicon encodes quantitative regulatory logic. Among the 215 CEs with non-additive activity, 190 showed synergistic activation and 25 showed attenuating or antagonistic effects. AND-like CEs capture classical combinatorial activation, exemplified by EICE^11^, NFAT::AP-1^13,14^ and novel configurations such as RUNX::CREB, whereas counteracting CEs implement AND-NOT-like or attenuating behavior that constrains activity despite an activating motif. Rare XOR-like CEs, in which two individually active motifs attenuate net output, suggest that composite pairing can encode mutual interference, a compact mechanism for enforcing mutually exclusive programs or initiating cell-state bifurcations. The enrichment of many functional CEs within TF co-bound regions supports their use at endogenous loci, and similarities to CAP-SELEX motifs provide complementary support for cooperative binding by TFs *in vitro*. Thus, composite elements act as context-encoding operators that disambiguate redundant simple motifs and increase the effective information content of regulatory DNA.

A notable finding is the diversity of AP-1-containing CEs. AP-1 has long been recognized as a central cooperative partner, including within the IFN-β enhanceosome^41^ and in the classical AICE^12^ and NFAT::AP-1^14^ elements. The advance of our study is therefore not the discovery of AP-1 cooperativity per se, but the systematic expansion of the AP-1-centered CE lexicon. The SMAD::AP-1, NFAT::AP-1 and ZEB::AP-1 CEs predict integration of AP-1 activity with TGF-β, Ca²⁺/calcineurin and Wnt pathways, respectively, illustrating how CE grammar encodes multi-pathway signal integration within a single composite unit.

GRACE provides both functional generalization and mechanistic interpretation of CEs. It learns composite grammar at single-nucleotide resolution from MPRA transcriptional readouts, yields interpretable contribution scores (CWMs) concordant with CEseek PWMs, and extends inference beyond the finite MPRA design to untested variants and configurations. Its convergence with ChromBPNet indicates that much of the composite lexicon is shared across modalities, whereas informative divergences distinguish mechanisms associated predominantly with chromatin accessibility from those associated with transcriptional activity. AlphaGenome further extends this perspective by enabling the simultaneous assessment of CE contributions across multiple chromatin and transcriptional features and across diverse tissue contexts. This joint-modeling perspective enables prioritization of motifs, CEs and variants predicted to alter chromatin state versus those primarily altering transcription within accessible chromatin.

Strikingly, ~30% of autoimmune-associated functional variants with motif-level GRACE interpretations mapped to CEs, underscoring the importance of composite grammar for interpreting non-coding variation in disease. CE-aware annotation matters for non-coding variation because CEs carry higher information content than their constituent motifs considered independently; variants disrupting composite grammar can thus have greater impact and evade single-motif scanning. Disruption of one motif collaterally affected its partner, consistent with CEs behaving as single functional units, while gain-of-function variants created *de novo* configurations. Thus, the framework captures both loss- and gain-of-function mechanisms. Coupling the CE lexicon with models that score transcriptional and chromatin consequences in a cell-context-aware manner should enable mechanistic stratification of regulatory variants and improve interpretation of GWAS loci.

CEseek identified two classes of motif pairs: “digital” pairings with ≤6 enriched configurations (the focus of this study) and a “fuzzy” subset with broad configurational tolerance. The fuzzy pairings may reflect mechanisms beyond a strict stereospecific model such as cofactor bridging, nucleosome positioning, or higher-order assemblies, and likely encode a softer, alternate layer of combinatorial control. Integrating the fuzzy regime with the digital lexicon, for example by tiling variable spacings in MPRA designs and linking to TF and co-factor perturbations, should help distinguish syntax imposed by direct TF-TF geometry versus chromatin- or complex-mediated interactions. More broadly, the CE lexicon will facilitate rational design of synthetic regulatory elements: selecting configurations with defined logic and actionable modality (accessibility vs. transcription) could yield enhancers with tunable, context-selective behavior for synthetic biology and therapeutic gene control.

Several limitations define important directions for future work. The low dynamic range of the minimal promoter used in the MPRA impaired the detection of repressing CEs, while the absence of measurable activity for many cCEs likely reflects the limited number of cellular contexts assayed. Next-generation libraries using alternative promoters assayed across a wider range of contexts should further expand CE discovery. Though cCEs were identified in the murine genome, their extensive characterization in human cell lines and genomic sequences indicates that the underlying composite grammar is largely conserved between mouse and human. Because the present study defines CEs primarily in motif rather than protein space, assigning specific TF paralogs to individual CEs will require complementary measurements of TF expression, occupancy, and perturbation. Future *in vivo* validation will benefit from approaches that directly assay TF co-occupancy at the same locus, including ChIP-ReChIP^42^ and SpDam-ID^43^, while perturbation of endogenous CEs and their cognate TFs will establish causal linkages and regulatory logic. Although the CE discovery herein focused on immune and hematopoietic OCRs, the CEseek-MPRA-GRACE framework is broadly applicable across cell types and tissues, and systematic expansion to additional tissues, developmental stages and signaling conditions will further enlarge the lexicon^44^. Community use of the accompanying Transcriptional Composite Element Portal, through which newly discovered CEs, model predictions, and future datasets can be integrated, will enable iterative refinement and expansion of the CE lexicon. Together, this work establishes composite elements as a major, functionally validated, and computationally accessible layer of the transcriptional regulatory code.

## RESOURCE AVAILABILITY

### Lead Contact

Further requests and information concerning this study should be addressed to the lead contact, Harinder Singh.

### Materials Availability

The TF-motif MPRA plasmid library is available upon request.

### Data and Code Availability

All sequencing data have been deposited into the IGVF Data Portal (https://data.igvf.org/): IGVFDS6431ZPLN, IGVFDS7446JCCY, IGVFDS5083RSLD, IGVFDS6230NHQE, IGVFDS2616NTKE, IGVFDS7386LSZJ, IGVFDS4765WKNZ, IGVFDS4645LTYC, IGVFDS2722MXQO, IGVFDS5743OUYJ, IGVFDS1360GZSW, IGVFDS6576ZAJI, IGVFDS9693DMOS, IGVFDS0273TRYL, IGVFDS2337RQZA, IGVFDS5855AVHR, IGVFDS9385FKEJ, IGVFDS2689EODI, IGVFDS5988JIIM, IGVFDS2955VGJT. Supplementary tables have been deposited on GitHub at https://github.com/Jingyu-Fan/TF_motif_lexicon/tables. The CEseek tool is publicly available at https://github.com/Jingyu-Fan/CEseek, and the GRACE model code is available at https://github.com/Jingyu-Fan/GRACE.

## ACKNOWLEDGEMENTS

This work was primarily supported by the National Institutes of Health (NIH) grant 5U01HG012041 (to N.S., J.D., and H.S.). This work was also funded by NIH grant RC2DK122376 (H.S.). Next-generation sequencing was performed by either the High-Throughput Genomics Core (UPMC) (formally the UPMC Genome Center) or the Health Sciences Sequencing Core (University of Pittsburgh). This research was supported in part by the University of Pittsburgh Center for Research Computing and Data, RRID:SCR_022735, through the resources provided. Specifically, this work used the HTC cluster, which is supported by the NIH award number S10OD028483. This work used bridge-2 (EM, RM, GPU) and ocean systems at the Pittsburgh Super Computing Center through the allocation of CIS240372 from the Advanced Cyberinfrastructure Coordination Ecosystem: Services & Support (ACCESS) program, which is supported by the U.S. National Science Foundation grants #2138259, #2138286, #2138307, #2137603, and #2138296. We thank Anshul Kundaje (Stanford), Lee Grimes and Nathan Salomonis (Cincinnati Children’s Hospital Medical Center) and Song (Stephen) Yi (Baylor) for valuable discussions and feedback.

## AUTHOR CONTRIBUTIONS

**Conceptualization:** J.F., V.K.C., H.S.

**Data Curation:** J.F.

**Formal Analysis:** J.F., V.K.C. (CEseek analysis of ImmGen ATAC-seq data)

**Methodology:** J.F., C.A.M., H.S.

**Project Administration:** J.F., N.A.P., N.S., H.S.

**Software:** J.F. (Python implementation of CEseek and development of GRACE), V.K.C. (original R prototype of CEseek)

**Investigation/ Data Generation:** J.F. (CEseek analysis, MPRA design and data analysis, GRACE model development and analysis), D.B. (MPRA assays), N.A.P. (Management and sequencing of MPRA libraries), D.R.H., V.F.Y., C.H.H., J.P.R. (MPRA training), S.K., R.T., N.S. (MPRA library construction).

**Visualization:** J.F.

**Resources:** N.S., H.S.

**Writing – Original Draft:** J.F., H.S.

**Writing – Review & Editing:** J.F., V.K.C., D.B., N.A.P., D.R.H., P.G., V.F.Y., S.K., J.D., J.P.R., R.T., C.A.M., N.S., H.S.

**Supervision:** C.A.M., N.S., H.S.

**Funding Acquisition:** H.S., N.S., J.D.

J.F. led the study; he re-implemented CEseek in Python, carried out various CEseek analyses, designed the MPRA library, performed all MPRA data analyses, developed the GRACE framework and performed GRACE analyses, curated and deposited the datasets, generated figures, and drafted the initial manuscript after its logical structuring with H.S..

V.K.C. created the original R prototype of CEseek and produced the initial cCE predictions using the ImmGen dataset and reported in BioRxiv^45^.

D.B., N.A.P., D.R.H., S.K., J.P.R., R.T., N.S., and H.S. were involved in the design, construction and characterization of the TF-motif MPRA library and generation of the experimental data involving transient transfections in GM12878, K562 and Jurkat cell lines.

N.A.P., V.F.Y generated the ATAC-seq data in GM12878 cells.

P.G. trained ChromBPNet models in K562 and GM12878 cells and generated contribution scores on OCRs.

C.A.M. guided the development of GRACE, refined the overall methodology together with

J.F. and H.S., and provided critical technical guidance and resources.

H.S. guided the scientific direction, secured funding, and oversaw all phases of the project.

J.F. and H.S. wrote the original draft.

V.C., D.B., N.A.P., D.R.H., P.G., V.F.Y., S.K., J.D., J.P.R., R.T., C.A.M., N.S., reviewed the manuscript. J.F. and H.S. made all revisions based on suggested changes.

All the authors have read and approved the final version of the manuscript and agree to be accountable for all aspects of the work.

## SUPPLEMENTAL INFORMATION

**Supplementary Table 1.** Position weight matrices (PWMs) of the 153 curated simple TF motifs and 5,063 cCEs identified in this study, related to **Fig. 1**.

**Supplementary Table 2.** cCE sequences (30-mers) identified by CEseek and incorporated into the TF-motif MPRA library design, including the number of stereospecific configurations tested per motif pair, related to **Figs. 1** and **2**. Simple motifs sequences used as qualitative references are also included.

**Supplementary Table 3.** Sequence-level transcriptional activities (log2FC) of 30-mer oligonucleotides in the TF-motif MPRA library, assayed in K562, GM12878, and Jurkat cells, related to **Fig. 2**.

**Supplementary Table 4.** Transcriptional activities and stereospecific configuration comparisons of cCEs in K562, GM12878, and Jurkat cells, including identification of configuration-dependent activity differences across motif pairs, related to **Fig. 2**. Activities of simple motifs used as qualitative references are included.

**Supplementary Table 5.** Non-additive transcriptional activities and regulatory logic classifications (AND, AND-NOT, XOR-like) of CEs in K562, GM12878, and Jurkat cells, related to **Fig. 2**.

**Supplementary Table 6.** Motif similarity report for functionally validated CEs compared with CAP-SELEX cooperative TF binding motifs, related to **Fig. 3**.

**Supplementary Table 7.** TF ChIP-seq datasets from K562 and GM12878 cells (CistromeDB) used for CE TF co-binding validation, related to **Fig. 3**.

**Supplementary Table 8.** CEseek-based enrichment of CE configurations in co-bound versus singly bound TF ChIP-seq peak regions (K562 and GM12878 cells), related to **Fig. 3**.

**Supplementary Table 9.** Motif similarity report for functionally validated CEs compared with *de novo* motifs from CistromeDB TF ChIP-seq datasets (K562 and GM12878 cells), related to **Fig. 3**.

**Supplementary Table 10.** CWM-PWM similarity scores (TomTom FDR) for activating cCEs in K562, GM12878, and Jurkat cells, related to **Fig. 4**.

**Supplementary Table 11.** K-means cluster assignments of OCRs based on GRACE-predicted transcriptional activity, with GRO-cap signal concordance at promoters and enhancers in GM12878 cells, related to **Fig. 4**.

**Supplementary Table 12.** GRACE-derived transcriptional activity and ChromBPNet-derived chromatin accessibility contribution scores for JASPAR 2024 CORE vertebrate motifs and functionally validated CEs, related to **Fig. 4**.

**Supplementary Table 13.** GRACE-predicted transcriptional activity and ChromBPNet-predicted chromatin accessibility for all 64 stereospecific configurations of validated CE motif pairs (K562 and GM12878 cells), including configurations not in the TF-motif MPRA design, related to **Fig. 5**.

**Supplementary Table 14.** GRACE-predicted allele-specific effects and CE/motif-aware interpretations for autoimmune disease-associated variants, including MPRA-validated emVARs (N=1,719) and extended predictions for 5,118 fine-mapped variants (PICS2 and Pan-UK Biobank, >30 diseases), related to **Fig. 6**.

**Supplementary Table 15.** Primers, probes, and library sequencing indices used for TF-motif MPRA library construction and sequencing, related to **Fig. 2**.

## METHODS

### CEseek computational framework

The CEseek pipeline was developed to systematically identify and analyze composite elements (CEs) within cis-regulatory DNA regions by leveraging position probability matrices (PPMs) to scan genomic sequences for transcription factor (TF) motif hits. Through pairwise motif scanning, CEseek evaluates combinations of all four motif orientations and user-defined spacing intervals, detecting motif co-occurrence patterns suggestive of potential cooperative or antagonistic interactions. Motif hits were identified using a PWM score threshold of at least 80% of the maximum possible score for the corresponding motif. Only motif hits exceeding this threshold were considered for subsequent CE analysis. Aligned motif hits, including spacer regions, are extracted and used to construct PPMs for the resulting CEs. By quantifying all possible binary configurations for user-defined spacing intervals, CEseek identifies dominant configurations exhibiting orientation and spacing preferences. Statistical enrichment of CEs is assessed via Fisher’s exact test, which compares motif pair frequencies in a test set against those in an appropriate background set (dinucleotide-shuffled sequences or other genomic sequences). For each CE configuration, each OCR was treated as a binary observation, indicating the presence or absence of at least one motif pair with the specified orientation and spacing. Multiple occurrences of the same configuration within an OCR were counted only once. Fisher’s exact test was then applied to compare the numbers of OCRs containing the configuration in the foreground and background sequence set. A chi-squared test is also implemented as an alternative in CEseek. Comparisons show strong agreement between the two approaches, with a Spearman correlation of 0.99 between the significance rankings obtained from Fisher’s exact test and the chi-squared test.

CEseek was implemented using Python programming language and incorporates several specialized libraries, including PyTorch for efficient tensor operations and computations, pybedtools^1,2^ for genomic interval manipulation, and pandas for data management and processing. The integration of GPU acceleration through PyTorch substantially enhances computational performance during motif scanning and sequence analyses. The CEseek tool is publicly available at https://github.com/Jingyu-Fan/CEseek. An archived version implemented in R is accessible at https://github.com/viren-v/CEseek.

### Compilation of a core simple TF motif set

A comprehensive library of TF motifs was compiled for systematic CEseek analysis. Initial collections of mouse and human TF motifs were obtained from the CIS-BP^3^ database (Build 2.00). A similarity matrix for all PWMs was generated using the HOMER^4^ function compareMotifs.pl with the “-matrix” option. For TFs represented by multiple PWMs, a minimal spanning tree was constructed using the similarity matrix, with PWMs as nodes. The PWM with the highest number of connections was selected as representative, and redundant PWMs were removed. In cases of ties, the PWM with the highest information content was retained. For PWMs from different TFs with a similarity score >0.9, a single representative PWM (with the highest information content) was selected. To ensure coverage, the CIS-BP collection was supplemented with curated TF motifs from the HOMER^4^ vertebrate motif library (http://homer.ucsd.edu/homer/motif/HomerMotifDB/homerResults.html), representing core (monomeric) binding sequences derived from ChIP-seq data. The combined motif set was manually curated to remove highly similar entries, resulting in a final library of 153 non-redundant simple TF motifs (Supplementary Table 1). See **Supplementary Note 1** for comparison of this core simple motif set to JASPAR 2024 CORE Vertebrates collection.

### Delineation of candidate CEs in ImmGen cis-Regulatory atlas

Chromatin accessibility data generated by the ImmGen^5^ consortium, which includes ATAC-seq profiles for 86 immune and hematopoietic cell types/states along with 4 non-hematopoietic cell types, were utilized for systematic CEseek analysis. The dataset comprises 512,595 open chromatin regions (OCRs), each of which is extended on both sides, yielding standardized 180-base pair (bp) regulatory regions. Importantly, the ImmGen ATAC-seq dataset was preprocessed using standard pipelines that include repeat masking, thereby minimizing potential contributions of repetitive elements (e.g., LINEs) to motif co-occurrence signals. CEseek was used to scan these OCR sequences systematically, evaluating all possible pairwise motif combinations derived from the library of 153 core TF motifs, with spacer lengths ranging from −5 to +10 base pairs. Each OCR was subsequently assigned to specific cell types or states based on its accessibility, defined as peaks falling within the 95th percentile for that particular cell type or state relative to others in the ImmGen dataset. Fisher’s exact test was employed to identify CE configurations that were statistically enriched in cell type- or state-specific OCRs, and the remaining OCRs were used as a background set (enrichment significance threshold p < 1e-20). Additionally, an independent analysis employing scrambled sequences as a control further validated the prevalence of cCEs, identifying 22,698 significantly enriched cCEs (p < 1e-10). Finally, dominant configurations characterized by preferential spacing and orientation between paired motifs were selected (significant motif pairs with ≤6 enriched configurations out of a possible of 64), resulting in 5,063 high-confidence cCEs. Corresponding PPMs for these selected cCEs were subsequently generated using aligned motif hits identified across the entire ImmGen OCR dataset. See **Supplementary Note 2** for details.

### Design of TF-motif MPRA library

A Massively Parallel Reporter Assay (MPRA) library comprising simple TF motifs and CEs was designed to enable comprehensive, systematic evaluation of motif activities *in vivo*. This library consisted of 60,714 sequences, each 30 bp in length. Initially, the set of 5,063 cCEs was scanned across ImmGen ATAC-seq peaks, and the top 10 motif hits with the highest scores for each cCE were selected. These selected hits were subsequently extended to uniform length 30 bp sequences. The sequences assigned to multiple cCEs were removed, resulting in a refined set of 3,795 unique cCEs. For each of these CEs, the four highest-scoring sequences were retained, and three additional mutated variants were systematically generated: one variant with mutations introduced into the left motif, another with mutations in the right motif, and a third variant with mutations in both motifs. Mutations involved substitutions at two consecutive positions of highest information content, converting nucleotide A to C and T to G, and vice versa. This strategy allowed for precise quantification of individual motif contributions within cCEs. Additionally, the MPRA library included 153 core simple motifs recognized by vertebrate TFs. For each simple motif, genomic sequences precisely matching motif length were identified and extended with selected random sequences to prevent unintended creation of alternative motifs. Up to 20 distinct sequences were selected for each simple motif based on their highest motif scores, alongside corresponding mutated sequences with substitutions at two consecutive positions of maximal information content. As controls, the library also contained 1,000 randomly generated 30 bp sequences without any recognizable motifs, serving as baseline measurements of background activity. To generate the 1,000 random 30 bp control sequences, we employed an iterative filtering strategy. Specifically, each candidate 30-mer was randomly generated and then scanned against our curated set of 153 core TF motifs. Only sequences with no detectable motif matches were retained. This process was repeated until 1,000 motif-free sequences were collected. See **Supplementary Note 3** for details.

### Construction of TF-motif MPRA library

The TF motif library sequences were cloned into the pGL4:23:ΔlucΔxbaI vector as described previously^6^ with modifications. To create the mpraΔorf library, barcoded oligos were inserted into Sfi*I*-digested pGL4:23:ΔlucΔxbaI by Gibson assembly (NEB, E2611) via the use of 1.1 µg of oligos and 1 µg of digested vector in a 50 µL reaction mixture and incubated for 60 min at 50°C. The reaction mixture was purified with 1.2x SPRI and eluted in 20 µL of elution buffer. Test transformation was performed to determine library coverage via the electroporation of 1 µL of ligated vector into 50 µL of electrocompetent 10-beta *E. coli* (NEB, C3020K) (2kV, 200 ohm, 25 µF). Similarly, for the scaled-up transformation, 1 µL of the library was transformed into 50 µL of electrocompetent cells. The electroporated cells were recovered in 950 µL of SOC media and then split into ten 1 mL aliquots of SOC. These cultures were subsequently recovered for an hour at 37°C, after which each culture was individually expanded in 20 mL of LB supplemented with 100 μg/mL carbenicillin (Sigma-Aldrich, C1389) in a shaker at 37°C for 6.5 hours. After incubation, all the cultures were pooled, and plasmid isolation was performed (Qiagen, 12963). To validate library complexity and connect barcodes to CE sequences, Illumina libraries were prepared for the mpra:Δorf library as described previously^7^ and sequenced via 2x150 PE reads on the Illumina NovaSeq platform with 15% PhiX spike-in. To create the final mpra:gfp library, 10 µg of mpra:Δorf library plasmid was linearized with 100 units of Asi*SI* (NEB, R0630) in a final 400 µL reaction and incubated overnight at 37°C. The linearized product was then column purified (Qiagen, 28104) and eluted in 30 µL. The GFP amplicon was amplified from pMPRAv3:minP-GFP via Q5 NEBNEXT Hot-Start (NEB M0494), 0.5 µM primer 200 and 0.5 µM primer 201 (Supplementary Table 15), following cycle conditions of 98°C for 30 s, 20 cycles (98°C for 10 s, 60°C for 15 s, 72°C for 45 s), and 72°C for 5 min. The amplified product was incubated with Dpn1 for 30 min at 37°C, followed by 0.5x reverse SPRI and 1.5x forward SPRI purification and elution in 40 µL EB. A second PCR was then performed using the 1:100 diluted purified PCR1 GFP amplicon as PCR1 and then column purified. This amplicon containing a minimal promoter, a GFP open reading frame and a partial 3’ UTR was then inserted by Gibson assembly using 1.6 µg of Asi*SI*-linearized mpraΔorf plasmid and 5.28 µg of the GFP amplicon in a 400 µL reaction for 90 min at 50°C, followed by 1.5x SPRI purification. The total recovered volume was redigested to remove the remaining uncut vectors by incubation with 50 units of Asi*SI*, 5 units of RecBCD (NEB, M0345), 10 µg of BSA, 1 mM ATP, and 1x NEB Buffer 4 in a 100-µL reaction mixture incubated overnight at 37°C, followed by 1.5x SPRI purification and elution with 40 µL of EB. To generate the final transfection-ready MPRA library, a test transformation was first performed using 1 µL of the final eluted mpra:gfp plasmid library and 50 µL of 10-beta cells to determine the library coverage. Finally, for scale-up transformation, 8 µL of mpra:gfp plasmid was electroporated (2kV, 200 ohm, 25 µF) into 200 µL of 10-beta cells. The electroporated bacteria were recovered in 12 mL of SOC, split across 6, 2 mL aliquots and incubated for 1 hour at 37°C. Then, each 2 mL culture was expanded to 500 mL of LB with 100 µg/mL carbenicillin and incubated for 16 hours at 37°C, followed by plasmid isolation via the Qiagen Gigaprep Kit (Qiagen, 12991).

### Transient transfections of TF-motif library in human cell lines

GM12878 cells (Coriell Institute for Medical Research) were cultured in RPMI medium (Corning, 10-040-CV) supplemented with 15% FBS (Gibco A52568-01), 1% GlutaMAX (Gibco 35050-061) and 1% Pen-Strep (Corning, 30-002-CI). For each transfection, 1*10^7^ cells were mixed with 10 μg of the TF-motif MPRA library in 100 μl of RPMI. These cells were transfected with a Neon transfection system (Thermo Fisher Scientific, MPK5000) using a kit (MPK10096B) with 3 pulses of 1200 V for 20 ms. A total of five biological replicates were performed, each using cells at a density of ~1 million cells/ml for the transfections. For each replicate, a total of 1.5*10^8^ cells were pelleted at *200 × g* and resuspended in 1.5 ml of RPMI medium containing 150 μg of the TF motif library. After transfection, each replicate was recovered at a density of 5*10^5^ cells/ml in 300 ml of RPMI medium containing 15% FBS, 1% GlutaMAX and 1% Pen-Strep. After 24 hours, the cells were pelleted at 200 × g and washed once with PBS. The cells were resuspended in 15 ml of RLT buffer provided with Qiagen Maxi RNeasy (Qiagen, 75162) and 300 mM DTT (G-Biosciences, 786227). The cells were then homogenized with a tissue homogenizer, TH (OMNI international), for 1 min at maximum speed and the lysates stored at –80°C. Jurkat T cells were cultured in RPMI medium (Gibco 61870-036) supplemented with 10% FBS (Gibco A52568-01) and 1% Pen-Strep (Corning, 30-002-CI). All other steps were the same as for GM12878 cells with the exception that the transfections using the Neon system involved 3 pulses of 1550 V for 10 ms.

K562 cells (CIMR) were cultured in Iscove’s modified Dulbecco’s medium (Corning 10-016-CV) supplemented with 10% FBS and 1% Pen-Strep. For each transfection, 1*10^7^ cells were mixed with 5 μg of the MPRA library in 100 μl of RPMI and transfected with the Neon transfection system using a kit with three pulses of 1450 V for 10 ms. Five biological replicates were performed as with GM12878 cells. For each replicate, a total of 1.5*10^8^ cells were pelleted at *200 × g* and resuspended in 1.5 ml of buffer R provided in the Neon Transfection Kit containing 150 μg of the TF motif library. After transfection, each replicate was recovered at a density of 5*10^5^ cells/ml in 300 ml of RPMI medium containing 15% FBS and 1% Pen-Strep. After 24 hours, the cells were pelleted at 200 × g and washed once with PBS. The cells were resuspended in 15 ml of RLT buffer and 300 mM DTT. The cells were then homogenized for 1 min at maximum speed and lysates were stored at –80°C.

### RNA extraction and cDNA synthesis

Total RNA was isolated from cells via Qiagen Maxi RNeasy (Qiagen, 75162) following the manufacturer’s protocol, including on-column DNase digestion by RNase-free DNase (Qiagen 79254). The final purified RNA was treated with 5 µl of SUPERase-In (Thermo Fisher Scientific, AM2696). Another DNase treatment was performed on total RNA via the addition of 5 µL of Turbo DNase (Thermo Fisher Scientific, AM2238) in 750 µL of total volume for 1 hour at 37°C. The reaction was terminated by the addition of 7.5 µL of 10% SDS (Life Technologies, 15553--035) and 75 µL of 0.5 M EDTA (Thermo Fisher Scientific, AM9260G), followed by incubation at 70°C for 5 minutes. All the DNase-treated RNA was then subjected to GFP mRNA pulldown. For this reaction, 900 µL of 20X SSC mixture (Life Technologies, 15557-044), 1800 µL of formamide mixture (Sigma Aldrich, 75--12--7) and 2 µL of 100 µM biotin-labeled GFP probe mixture (GFP_BiotinCapture_1--3, Supplementary Table 15) were added, and the volume was adjusted to 3600 µL. The mixture was then incubated at 65°C for 2.5 hours with intermittent inverting of the tubes every 30 min. The biotin probes were captured via 400 µL of prewashed streptavidin beads (Thermo Fisher Scientific, 65002). The streptavidin beads were washed twice with Buffer-A containing 0.1 M NaOH (Sigma-Aldrich, S2770) and 0.05 M NaCl (Thermo Fisher Scientific, AM9760G) and once with Buffer-B containing 0.1 M NaCl. The beads were then eluted in 500 µL of 20X SSC and added to the RNA probe sample. The hybridized RNA-probe-bead mixture was mixed with a HulaMixer (Thermo Fisher Scientific, V.3A01) at room temperature for 15 minutes. The beads were captured with a DynaMag magnet (Thermo Fisher Scientific, 12321D) and washed once with 1x SSC and twice with 0.1x SSC. The extraction of RNA was performed by the addition of 25 µL of DEPC-treated water (Ambion, AM9906) and heating of the mixture for 2 minutes at 80°C, followed by immediate collection of the eluent on a magnet. A second elution was performed by incubating the beads with an additional 25 µL of water at 80°C. The eluted RNA was then processed for the final DNase treatment with 1 µL of Turbo DNase in a total of 56 µL of reaction. The mixture was incubated for 60 minutes at 37°C, followed by inactivation with 1 µL of 10% SDS. The final DNase-treated GFP mRNA was eluted via RNAClean XP SPRI beads (Beckman, A63987) in 35 µL of DEPC-treated water. First-strand cDNA was synthesized via the use of 30 µL of DNase-treated GFP mRNA with SuperScript III and primer 19 (Supplementary Table 15) according to the manufacturer’s recommended protocol. Single-stranded cDNA was purified via AMPure SPRI (Beckman, A63881) beads and eluted in 30 µL of EB.

### Generation and sequencing of Tag-seq libraries

To minimize amplification bias during the creation of cDNA tag sequencing libraries, samples were amplified via qRT-PCR to estimate the relative concentrations of GFP cDNA. qRT-PCR was performed via the use of a 1 µL cDNA sample in a 10 µL PCR mixture containing 5 µL of Q5 NEBNext Ultra II master mix, 1.7 µL of SYBR Green I diluted 1:10,000 (Life Technologies, S-7567) and 0.5 µM of TruSeq_Universal_Adapter and MPRA_Illumina_GFP_F primers (Supplementary Table 15). The samples were amplified with the following conditions: 98°C for 20 s; 40 cycles of 98°C for 10 s, 62°C for 15 s, and 72°C for 30 s; and 72°C for 2 min, followed by melt curve analysis. For sequencing of plasmid library, serial dilutions of the plasmid library were prepared from 1000 pg to 1 fg via 10-fold dilutions. A standard curve was plotted with the plasmid library dilutions and based on the threshold cycles for the cDNA samples, a plasmid dilution with the same cycle threshold value was used for adaptor ligation. To add Illumina sequencing adapters, cDNA samples and 5 plasmid library controls were diluted to normalize the cDNA replicate with the lowest concentration or highest threshold cycle, and 10 µL of normalized sample was amplified using the reaction conditions from the qRT-PCR scaled to 50 µL, without adding SYBR Green to the reaction and by using only n-1 amplification cycles (where n= the cycle number obtained from qRT-PCR). The amplified cDNA was 2x SPRI purified and eluted in 30 µL of EB.

A second PCR was performed to add the individual index primers to each sample. The PCR was performed with 20 µL of purified PCR 1 elute in a 50 µL Q5 NEBNext Ultra II reaction with 0.5 µM of TruSeq_Universal_Adapter primer and Illumina_Multiplex primer containing a unique 8 bp index for sample demultiplexing postsequencing (Supplementary Table 15). The samples were amplified at 98°C for 20 seconds, 6 cycles of 98°C for 10 sec, 62°C for 15 sec, and 72°C for 30 sec, and 72°C for 2 minutes. Indexed libraries were 2x SPRI purified and quantified on a Qubit Flex (Thermo Fisher Scientific, Q33326) using a dsDNA HS Assay Kit (Thermo Fisher Scientific, Q33230) and pooled according to molar estimates from TapeStation (Agilent Technologies, 4200 TapeStation) quantification. The samples were sequenced via 1x20 bp reads on the Illumina NextSeq 2000 platform, with 80 M reads per cDNA library replicate and 60 M reads per plasmid library replicate.

### MPRA data processing

To process the MPRA data, we employed an established pipeline (https://github.com/tewhey-lab/MPRA_oligo_barcode_pipeline). The barcode-oligo sequencing FASTQ files were parsed using *MPRAmatch* function to create the barcode-oligo dictionary, retaining only barcode-oligo links with exact 30 bp matches. Barcode count tables were subsequently generated using *MPRAcount* function.

To quantify the activities of simple motifs and CEs, the analysis was conducted at three levels: barcode, oligo, and motif. First, barcode-level count tables were processed with DESeq2^8^ to remove barcodes with excessively large counts inconsistent with those of the biological replicates. This filtering used Cook’s distance, applying a 99% quantile threshold to exclude outliers. After filtering, the barcodes were aggregated into their corresponding oligos, and only those with at least 30 unique barcodes were retained for further analysis. The activities of the 30 bp oligos were assessed by performing DESeq2 differential analysis, which involved comparing the total barcode counts associated with each oligo in the RNA and DNA samples. Normalization and size factor estimation were performed using DESeq2’s internal methods. Additionally, cell type-specific dispersion values were calculated by analyzing only RNA replicates, and these refined dispersions were used to improve the accuracy of the final analysis. To further increase the reliability of the differential activity measurements, log fold change shrinkage was applied via the lfcShrink function from DESeq2, which helps reduce the impact of extreme log fold changes in oligos with low counts and provides more stable effect size estimates. To assess the activity of simple and composite TF motifs accurately, a comprehensive approach was employed. For simple motifs, activities were determined by comparing the RNA/DNA log2-fold change in oligos containing the simple motif (sequence activity) with matched mutated control oligos (motif activity). For composite motifs, sequence and motif activities were estimated similarly by comparing RNA/DNA ratios for oligos containing the CE to those in matched double-mutant control oligos. Additionally, the contributions of individual motifs (motif1 and motif2) within a CE were examined by comparing the RNA/DNA ratios in oligos with one mutated motif to those in matched double-mutant control oligos. Synergistic effects between motifs within a CE were assessed by comparing the motif activity of the CE to the sum of the contributions from motif 1 and motif 2.

Simple and composite motifs were classified as activating or repressing based on both sequence and motif activity measures. Specifically, a motif was considered activating or repressing if both activities were significant (false discovery rate, FDR < 0.05) and exhibited the same directionality (activating or repressing). Synergistic effects were considered statistically significant when the p-value was < 0.05. See **Supplementary Note 5** for details.

### Comparison of cCEs with motifs from TF ChIP-seq and CAP-SELEX datasets

To provide evidence for TF binding *in vivo* and *in vitro,* the cCEs identified by CEseek were compared with motifs obtained from chromatin immunoprecipitation sequencing (ChIP-seq) and CAP-SELEX datasets. Specifically, a comprehensive collection of uniformly processed TF ChIP-seq datasets from CistromeDB^9–11^ was assembled for hematopoietic (K562) and B cell (GM12878) cell lines. This dataset comprised 962 ChIP-seq experiments corresponding to 337 unique TFs in K562 cells and 370 experiments representing 160 unique TFs in GM12878 cells. From these datasets, a total of 14,897 *de novo* motifs were identified. Subsequently, each cCE identified by CEseek was systematically compared against this database of *de novo* motifs using the Tomtom^12^ motif comparison tool to assess motif similarity. Additionally, a similar comparative analysis was conducted between cCEs and a set of 1,749 (618 + 1,131) CAP-SELEX-derived heterodimeric motifs^13^. In this case, each CAP-SELEX motif was compared against the entire database of cCE motifs using the same computational approach (Tomtom). See **Supplementary Note 8** for details.

### Enrichment analysis of CEs in TF ChIP-seq datasets

To determine whether CEs are specifically associated with co-binding of cognate TFs *in vivo*, a three-step analysis pipeline was implemented. First, simple TF motifs within each CE were matched with molecularly relevant ChIP-seq datasets. A simple motif was considered matched to a TF ChIP-seq dataset (i) if the TF targeted by the ChIP-seq experiment belonged to the same TF family as the simple motif or (ii) if the simple motif significantly matched one of the top 10 motifs from the TF ChIP-seq dataset (FDR < 0.05, assessed using Tomtom). The less stringent *in vivo* TF binding evidence (ii) to CE regions is reported separately in Supplementary Table 8. Second, all CEs exhibiting significant transcriptional activity (activating or repressing) and synergistic effects were evaluated for enrichment within peaks from matched TF ChIP-seq datasets. This enrichment analysis was conducted using CEseek, with dinucleotide-shuffled sequences (four times the number of test sequences) serving as the background. Statistical significance was assessed using Fisher’s exact test, with a stringent enrichment threshold of p ≤ 1e-10. Finally, a selected set of CEs was assessed for enrichment specifically within regions co-bound by both relevant TFs (by at least 1 bp overlap). This analysis was conducted by comparing the occurrence of the CE within co-bound ChIP-seq peak regions to its occurrence within singly bound regions as the background, employing CEseek and Fisher’s exact tests for statistical evaluation. See **Supplementary Note 8** for details.

### GRACE model

To predict cell type-specific regulatory activities in short DNA sequences, we developed a predictive model named GRACE (Genome Regulatory Analysis of Simple and Composite Transcriptional Elements). The GRACE model architecture starts with a convolutional layer consisting of multiple convolution blocks applied to DNA sequences that are transformed into one-hot encoded matrices. These blocks use different kernel sizes to capture motifs of varying lengths. This is followed by a transformer-based attention mechanism that analyzes motif interactions. The convolutional layer helps identify transcription factor (TF) binding sites of various sizes, whereas the self-attention mechanism, which is enhanced with positional encoding, captures the complex interactions between these motifs. This combination allows the model to effectively represent and analyze intricate regulatory elements within sequences.

To mitigate overfitting and enhance the model’s generalizability, we employed dropout layers and data augmentation techniques, such as random sequence additions and reverse complement usage, to diversify the training data. Additionally, early stopping was used to halt training once validation loss failed to improve for a certain number of epochs, ensuring optimal model performance without overfitting.

The GRACE model was implemented via PyTorch and PyTorch Lightning, with a custom DataModule that handles data loading, splitting, augmentation, and normalization of activity values, ensuring streamlined and efficient data handling. The model was trained on MPRA experimental datasets containing oligo activities, which were preprocessed and divided into training, validation, and test sets. We assigned variant sequences that were derived from a given parent sequence to the same subset, preventing data leakage, thereby enhancing the model’s ability to generalize predictions to all possible single nucleotide variants. Preprocessing involves filtering sequences by barcode count, augmenting them with random segments to mitigate positional bias, and incorporating reverse complement sequences to improve robustness and generalizability. Sequence padding was applied to ensure uniform input lengths for efficient batch processing. The model’s attention mechanism was restricted to 30 bp blocks, allowing the model to focus on local regulatory features while maintaining computational efficiency and avoiding artificial long-distance attention created by random sequence augmentation.

Training was conducted via the AdamW optimizer, which incorporates weight decay to prevent overfitting, along with a learning rate scheduler to dynamically adjust the learning rate throughout training, improving convergence. Early stopping was also employed to prevent overfitting by monitoring the validation loss and halting training once the performance ceased to improve. The evaluation metrics included the mean squared error (MSE), Spearman’s correlation, and Pearson’s correlation, providing a comprehensive assessment of the model’s ability to predict regulatory activity. To validate the model, predictions on withheld test sequences were compared directly to experimentally measured activities, ensuring reliable model performance. See **Supplementary Note 10** for the details for GRACE model training and **Supplementary Note 11** for the benchmark of GRACE model with other methods.

The code for the GRACE model can be found at https://github.com/Jingyu-Fan/GRACE.

### GRACE contribution weight matrix generation and comparison with CEseek PWMs

To elucidate the motifs contributing to transcriptional activity, in silico saturation mutagenesis was performed, systematically mutating each nucleotide position within the input DNA sequences to evaluate their individual contributions to transcriptional activity. For each 30-bp oligonucleotide, single-base substitutions were introduced at every position, and the transcriptional activities (log2-fold changes) of these variants were predicted using GRACE. The predictions were organized into 4 × L matrices (rows corresponding to nucleotides A, C, G, and T; columns corresponding to sequence positions). Contribution weight matrices (CWMs) were then generated by subtracting the mean log2 activity of each position from the activity of each nucleotide variant at that position. This normalization step removed position-specific biases. To produce a sparse representation highlighting key nucleotide contributions, only the scores corresponding to nucleotides present in the original (wildtype) sequences were retained, with all other entries set to zero. The resulting matrices were visualized as contribution scores.

To compare these GRACE-derived CWMs systematically with CEseek-generated PWMs, the CWMs were first converted into position probability matrices (PPMs) via the Softmax function, ensuring that the probabilities at each position were summed to 1. To assess similarity, each resulting PPM was compared to its corresponding CEseek-derived PWM via Tomtom. For robust statistical comparison, 1,000 randomized background PWMs were generated for each PWM by shuffling positions and nucleotides. The similarity between each CWM-derived PPM and its corresponding CEseek-derived PWM, as well as the background PWMs, was evaluated, and the false discovery rate (FDR) was calculated. Multiple oligonucleotides representing each CE were analyzed, and the median of the negative log10-transformed FDR values was computed. A threshold of FDR < 0.001 was employed to define significant similarity between a CE’s GRACE-derived CWM and its corresponding CEseek-derived PWM.

### GRACE predictions of transcriptional activity of open chromatin regions

The GRACE model was applied to predict the transcriptional activities of the OCRs. Because GRACE requires 50 bp sequences as inputs and OCRs typically exceed this length, a stepwise procedure was implemented. First, each OCR was segmented into overlapping 50 bp subsequences using a 1 bp sliding window, and the activity (log2-fold change, log2FC) of each subsequence was predicted using GRACE. All the predicted activities for the 50-mer sequences within each OCR were subsequently ranked from highest to lowest. The subsequence with the highest predicted activity was selected and included in the overall activity score of the OCR if it exceeded a predefined threshold (log2FC = 0.2). This process was iteratively repeated, excluding subsequences overlapping previously selected segments, until no remaining 50-mer sequences surpassed the activity threshold. The sum of the selected subsequences’ activities represented the final GRACE-predicted transcriptional activity of the OCR. The H3K27ac ChIP-seq signal of an OCR is calculated as the average read per million (RPM) for the 1 kb size window centered at the OCR and the ATAC-seq signal.

### Systematic analysis of contributions of simple motifs and CEs to genomic features

The contributions of simple motifs and CEs to transcriptional activity and chromatin accessibility were systematically evaluated using genome-wide contribution scores derived from the GRACE and ChromBPNet models, respectively. AlphaGenome was additionally used to evaluate the contributions of simple and composite motifs to nascent transcription and histone modifications; due to bandwidth limitations, these analyses were performed on a sampled subset of genomic regions (n=100). For this analysis, the collection of simple motifs included 879 motifs sourced from the JASPAR 2024 vertebrate core collection^14^. The CE collection consisted of 951 unique CEs identified as being active in the MPRA experiments conducted in the K562, GM12878, and Jurkat cell lines. To identify instances of simple and composite motif hits within OCRs, CEseek was employed. The contribution scores obtained from GRACE and ChromBPNet were organized into CWMs, which were formatted as 4 × L matrices (rows representing nucleotides A, C, G, and T; columns representing sequence positions). For each motif occurrence, the contribution scores were summed and averaged by dividing by the number of motif hits. Each averaged CWM was subsequently normalized through elementwise multiplication with the corresponding PPM derived from CEseek-detected motif hits. The resulting sum was divided by the motif length to yield the motif’s normalized contribution score. The baseline contribution scores were computed similarly using ±50 bp flanking regions around motif hits. The final motif contribution was calculated as the fold difference between the motif hit region and the baseline flanking region. For visualization, the aggregated CWMs were computed over a window spanning the motif region extended by a fixed number of flanking bases (typically ±10 bp). Baseline contribution levels were estimated independently from the distal flanking regions (typically the outer ±50 bp relative to the motif center) by averaging contribution scores across these regions. These baseline values were used for visualization as reference levels (e.g., dashed lines in CWM logos) but did not alter the underlying CWM values.

### GRACE analysis of GWAS variants associated with autoimmune diseases

The GRACE model was employed to predict and interpret the functional impact of disease-associated genetic variants identified through genome-wide association studies (GWASs). A recently published MPRA dataset comprising functional assays conducted with 150 bp reference and variant sequences in an Epstein-Barr virus-transformed B-cell line derived from a lupus patient was used^15^. From this MPRA dataset, expression-modulating variants demonstrating ambiguous allele-specific effects without statistical significance (|log2FC| > 0.1 and FDR > 0.05) were excluded. A total of 1,719 expression-modulating variants were subsequently compiled, 319 of which exhibited significant allele-specific effects (|log2FC| > 0.6 and FDR < 0.05). To predict the variant effects, GRACE activity predictions were performed on 50 bp sequences centered on each variant site, comparing reference and alternative alleles. The difference in predicted activity between the two alleles was utilized as a measure of the variant effect. An absolute value of 0.2 of a GRACE-predicted variant effect was used as the threshold for determining significant variant effects. Applying this criterion resulted in the successful prediction of 68 functional variants. GRACE-generated contribution scores for these 50 bp sequences were then analyzed to interpret the TF motifs (simple or CE) underlying the observed variant effects.

To extend this analysis, we incorporated a larger set of autoimmune disease-associated variants curated from PICS2^16^ fine-mapped GWAS datasets and the Pan-UK Biobank^17^ (Pan-UKB) resource. Pan-UKB genome-wide association summary statistics were obtained from publicly available datasets and parsed to extract variant-level information, including genomic coordinates, alleles, effect sizes, and association statistics. Disease-specific datasets were selected based on phecode annotations corresponding to autoimmune conditions, including multiple sclerosis (MS), rheumatoid arthritis (RA), and systemic lupus erythematosus (SLE). In parallel, PICS2-derived variants were used to obtain a fine-mapped set of candidate causal SNPs across >30 autoimmune diseases, yielding 5,118 variants analyzed in this study (Supplementary Table 14). These variants were evaluated using GRACE models trained on TF-motif MPRA data in GM12878 B cells, and their predicted transcriptional effects were compiled as part of the “Variant impact contribution plots” in the CE portal.

---

**Extended Data Fig. 1.**
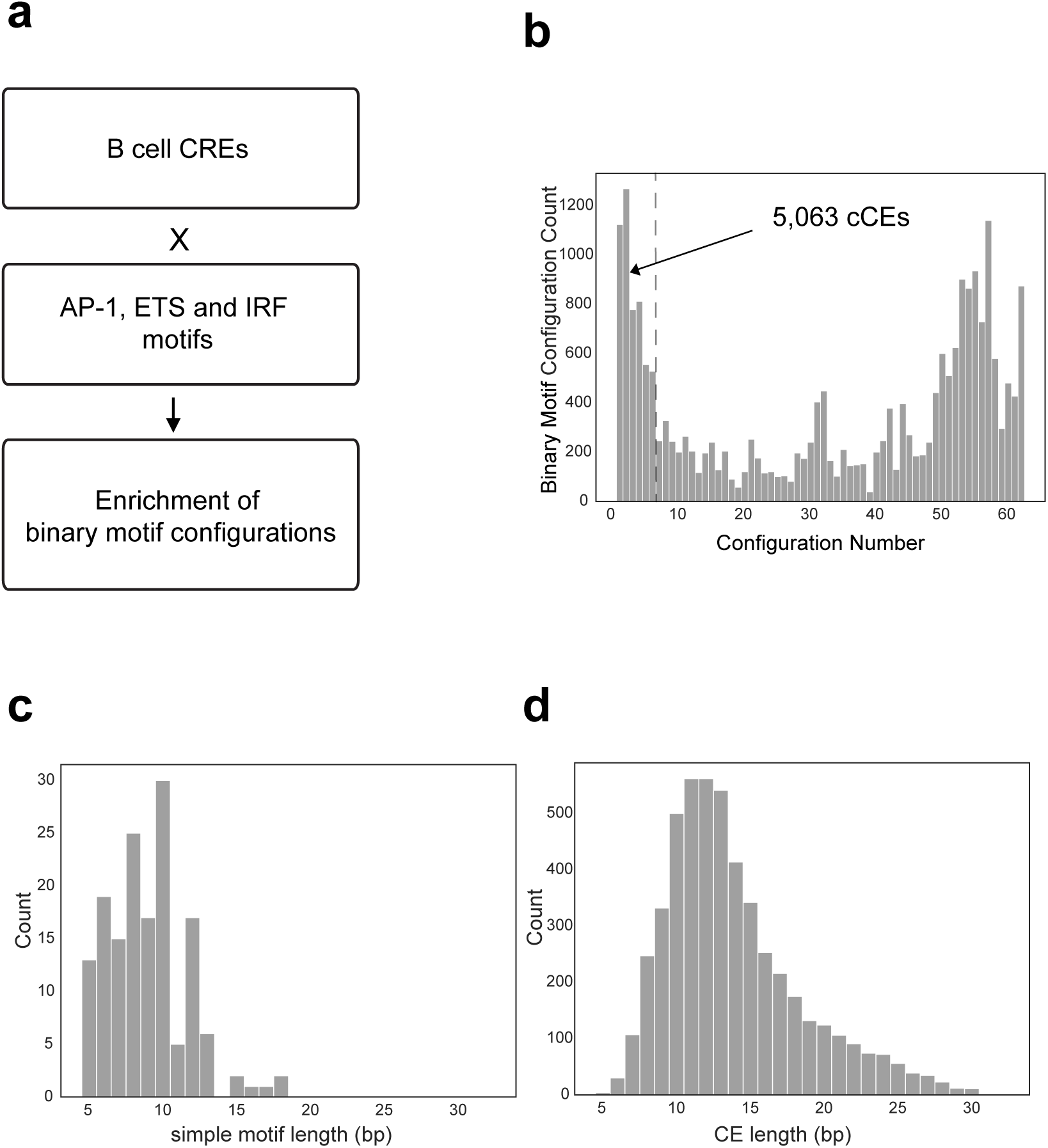
CEseek testing and cCE properties. **(a)** Proof-of-concept workflow using CEseek to recover known CEs composed of ETS, AP-1, and IRF motifs from murine B-cell enhancers characterized by STARR-seq. **(b)** Histogram displaying the distribution of binary motif pair configuration counts (n=22,698, see **Fig. 1d**) with their numbers of statistically enriched configurations (n=1 to 64). The left end of the distribution highlights motif pairs with ≤6 unique configurations out of a possible of 64 that were designated as cCEs (see **Fig. 1e**). The right end of the distribution highlights motif pairs with >50 statistically enriched configurations out of a possible of 64. **(c, d)** Histograms displaying the length distributions of 153 simple TF motifs **(c)** and 5,063 cCEs **(d)**.

**Extended Data Fig. 2.**
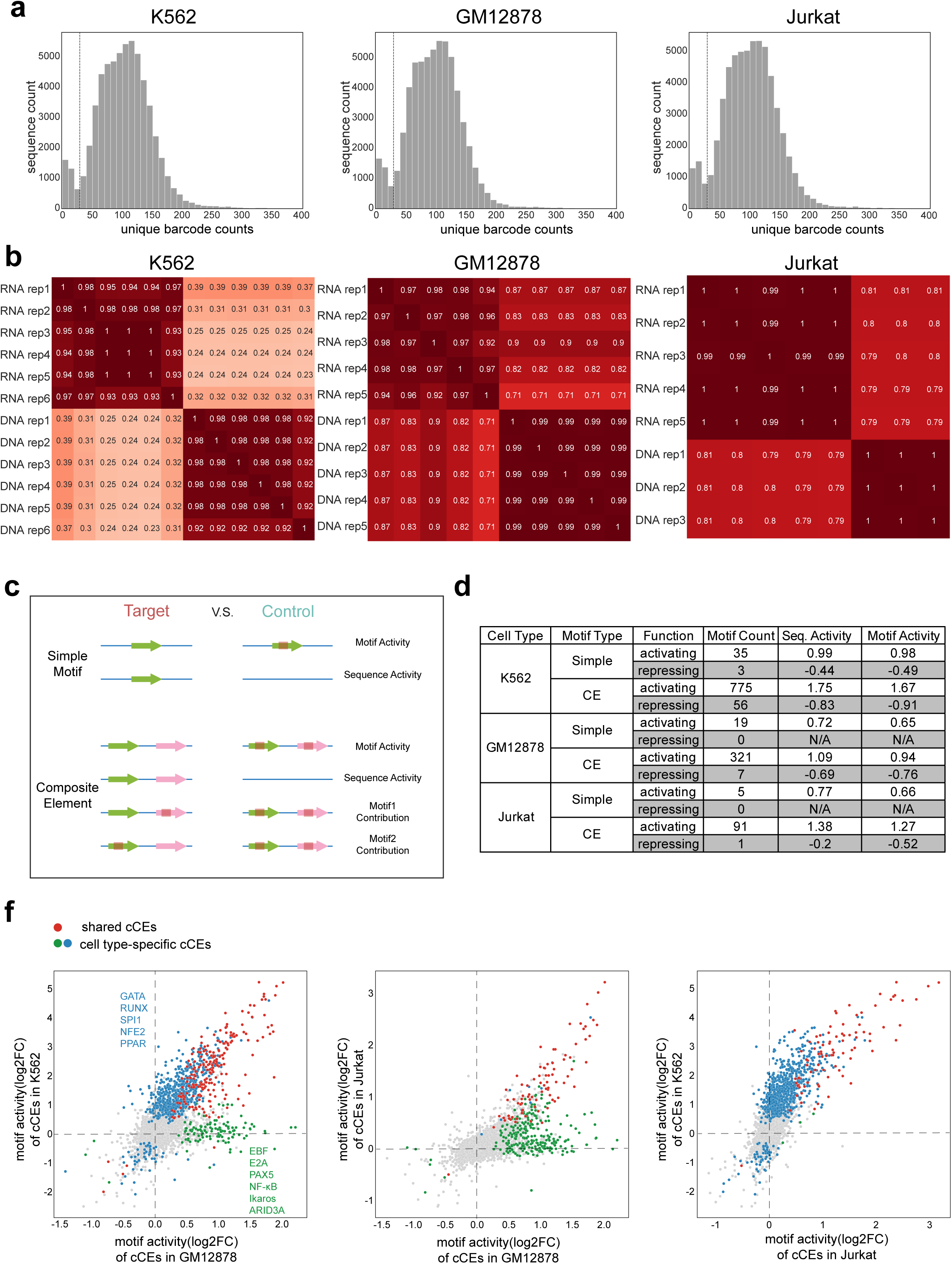
Analysis of cCEs using a custom TF motif MPRA library. **(a)** Barcode count distribution. Histograms showing the distribution of unique barcode counts recovered from RNA for TF motif library sequences after transfection in K562 (left), GM12878 (middle), and Jurkat cells (right). Sequences with ≥30 unique barcodes (dashed line) were used for downstream analyses. **(b)** Reproducibility of the biological replicates. Heatmap displaying pairwise Pearson correlation coefficients for aggregated barcode counts linked with a given TF motif sequence across RNA and DNA biological replicates. **(c)** Schematic illustrating the determination of simple and composite motif activities from the MPRA datasets. For CEs, motif activity was calculated using DEseq2 by comparing the RNA/DNA log2 fold change (log2FC) of motif-containing sequences (sequence activity) as test values and the RNA/DNA log2 fold change (log2FC) of matched double-mutant controls as reference values. To determine individual motif contributions within a CE, the RNA/DNA log2FC values of sequences carrying single-motif mutations were analyzed, with those from matched double-mutant sequences used as controls. For simple motifs, motif activity was similarly calculated by comparing motif-containing sequences (sequence activity) with matched mutated control sequences. **(d)** Numbers of functional simple and composite TF motifs revealed by MPRA assays. Table summarizing the counts of functional (activating or repressing) simple and composite motifs identified in K562, GM12878, and Jurkat cells, along with their median sequence and motif activities. **(e)** Comparisons of cCE activities across cell types. Scatter plots of cCE motif activity (log2FC) measured in GM12878 vs. K562 (left), GM12878 vs. Jurkat (middle), and Jurkat vs. K562 (right) cells. The green and blue dots indicate cell type-specific activating or repressing cCEs in the indicated cell types. The red dots are cCEs that are activating or repressing in both cell types. The gray dots are cCEs with no detectable activity in the indicated cell types. Representative simple TF motifs that are constituents of cell-type specific CEs are shown in the left panel.

**Extended Data Fig. 3.**
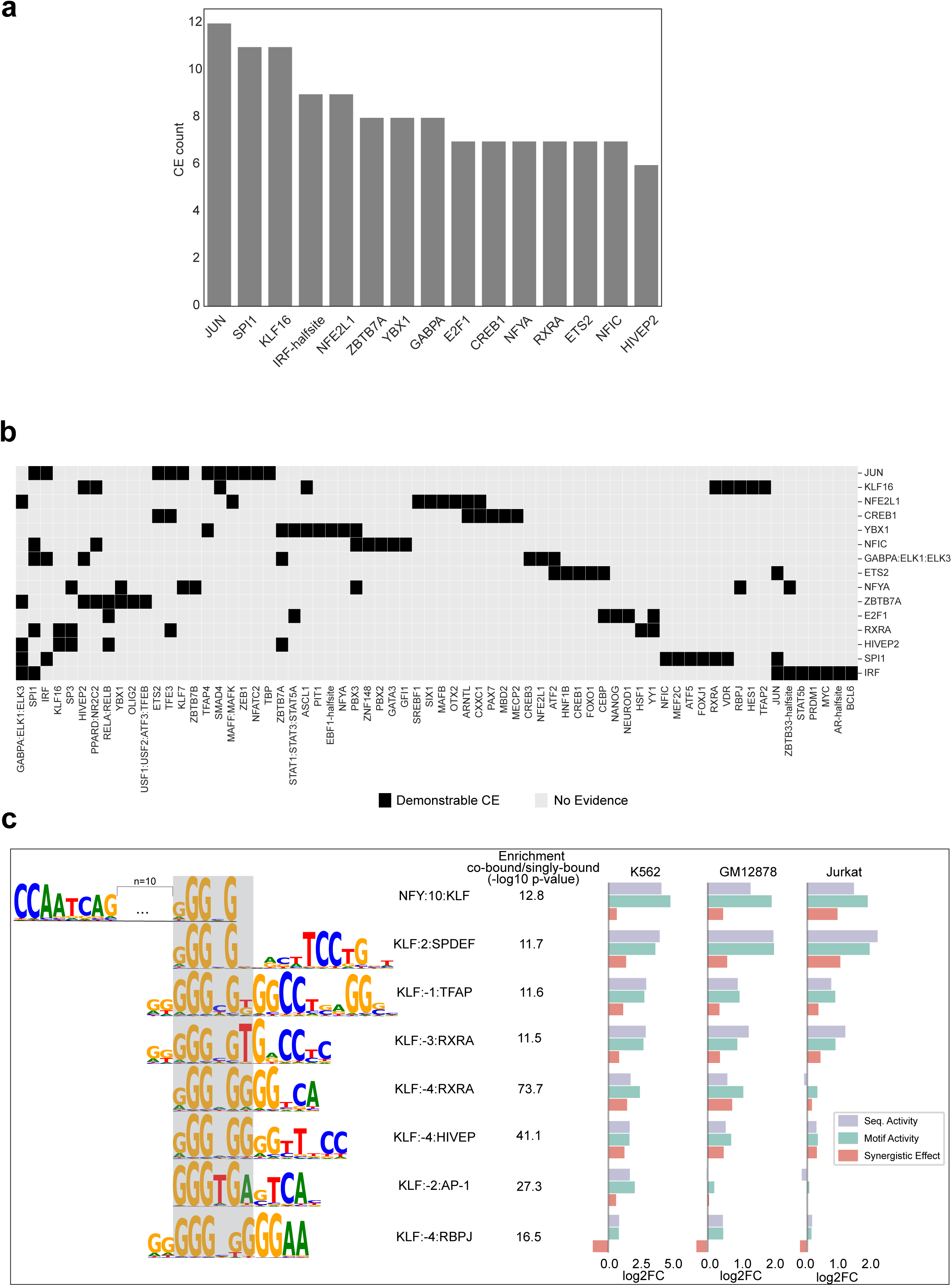
CEs are enriched in cognate TF-bound genomic regions. **(a)** Prevalent simple TF motifs contained within CEs. Bar plot showing the counts of simple TF motifs within CEs that are enriched in cognate ChIP-seq datasets (CistromeDB). **(b)** Diversity of CEs. Grid displaying the pairs of simple TF motifs that comprise experimentally demonstrable CEs. Rows are the top 15 simple TF motifs with the highest CE counts, displayed in **(a)**. Columns are the more diverse simple TF motifs that the prevalent set (**a**) is paired with. **(c)** Diversity of CEs comprising KLF motifs. Sequence logos of the KLF families of CEs are shown on the left. The bar plots show the measured sequence, motif, and synergistic activities of each CE in K562, GM12878, and Jurkat cells. Supporting evidence for TF co-binding was obtained from available CistromeDB ChIP-seq datasets for the specific TF family members listed in Supplementary Table 8.

**Extended Data Fig. 4.**
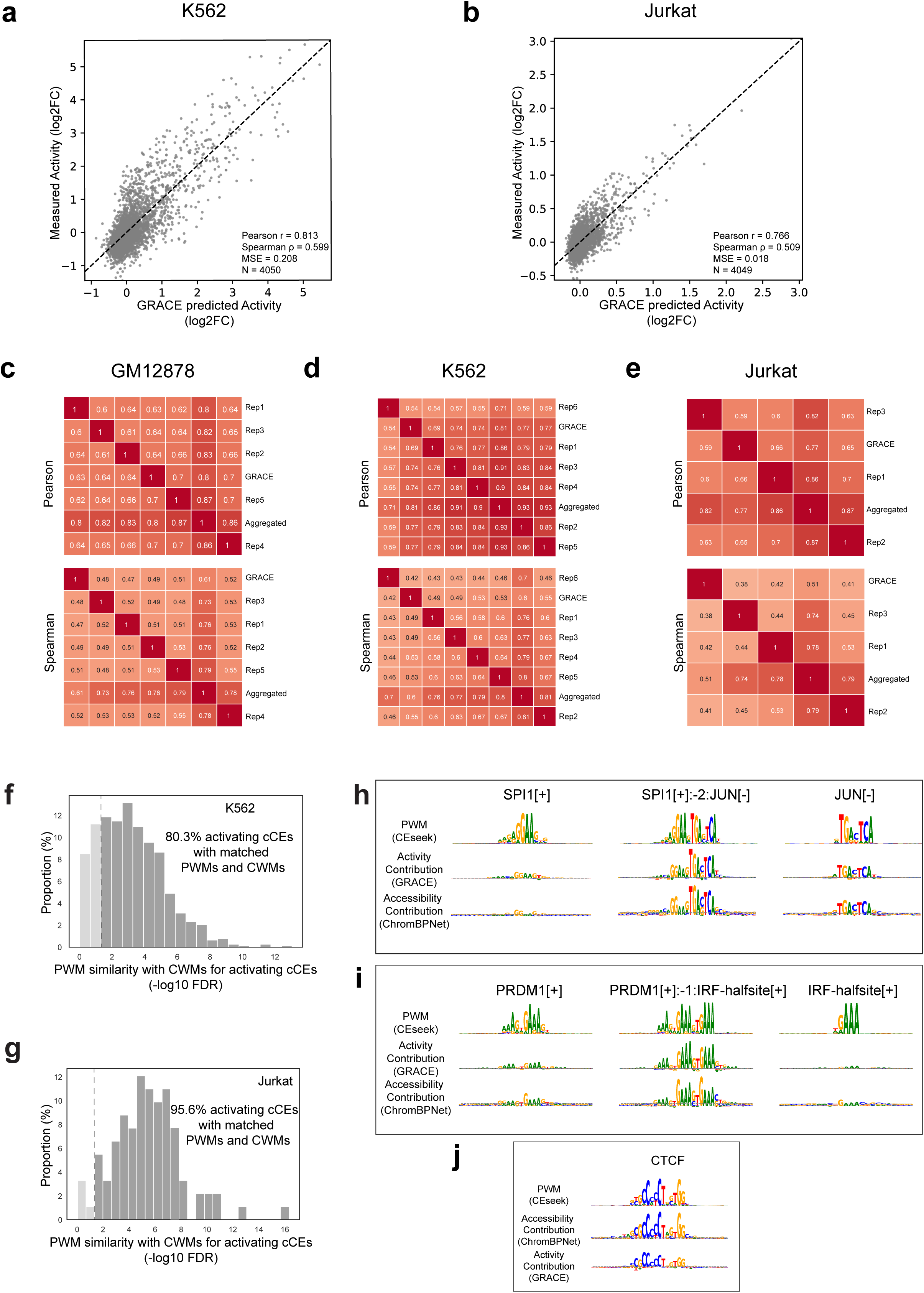
GRACE performance and CE contribution analysis. **(a, b)** Prediction accuracy of GRACE models trained on the K562 **(a)** and Jurkat **(b)** MPRA datasets, as shown in **Fig. 4b**. Scatter plots comparing GRACE-predicted transcriptional activities for withheld test sequences to experimentally measured MPRA activities (log2FC). Pearson and Spearman correlations are indicated. **(c-e)** Correlations of GRACE predictions with MPRA replicates. Heatmaps showing pairwise Pearson (**top**) and Spearman (**bottom**) correlations between GRACE predictions and MPRA measurements across individual biological replicates, as well as with the aggregated set of MPRAs in GM12878 **(c)**, K562 **(d)** and Jurkat cells **(e)**. **(f, g)** Histograms showing the false discovery rate (FDR) of the similarity between CEseek PWMs and GRACE CWMs for activating cCEs in K562 **(f)** and Jurkat cells **(g)**. The fraction of matching cCEs (FDR < 0.05) is indicated in each panel. **(h-i)** Examples of CEs and individual TF motifs with their contributions to chromatin accessibility and transcriptional activity, as in **Fig. 4g,h**. Examples of CEs: left and right columns show constituent simple motifs; the middle column shows the CE. The first row in each panel displays sequence logos (PWMs) based on CEseek. The second and third rows provide the CWMs for transcriptional activity (second row) and chromatin accessibility (third row) using GRACE and ChromBPNet models, respectively. The dashed gray lines indicate baseline contributions estimated for 50 bp flanking regions (see **Methods**). **(j)** Example of the CTCF motif, which was not included in the MPRA library but was identified by GRACE from related sequences; shown to illustrate that CTCF contributes more strongly to chromatin accessibility (ChromBPNet) than to transcriptional activity (GRACE).

**Extended Data Fig 5.**
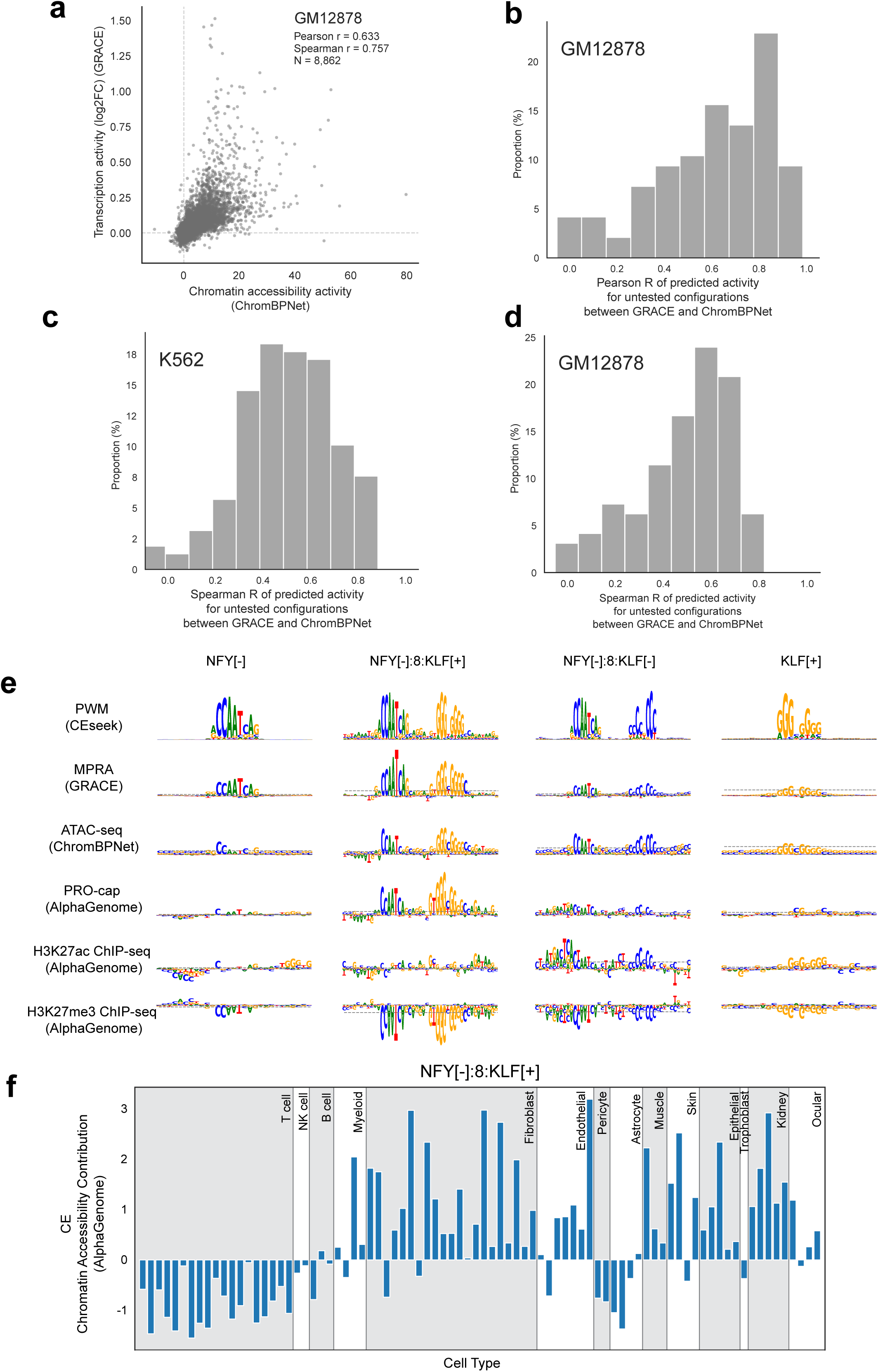
Multi-modal analysis of untested CE configurations. **(a)** Scatter plot comparing GRACE-predicted transcriptional activity and ChromBPNet-predicted chromatin accessibility for CE configurations not included in the TF-motif MPRA design (GM12878 cells). **(b-d)** Histograms of Pearson (**b**) or Spearman (**c,d**) correlation coefficients between GRACE and ChromBPNet predictions across individual motif pairs for untested configurations. (**b**) GM12878, Pearson R (median R = 0.651). (**c**) K562, Spearman R. (median R = 0.494) (**d**) GM12878, Spearman R (median R = 0.533). These complement **Fig. 5b**. **(e)** Example of NFY::KLF CEs and their constituent simple motifs, illustrating motif contributions across multiple regulatory modalities. Shown are position weight matrices (PWMs; CEseek) and model-derived contribution profiles (CWM) from GRACE (MPRA), ChromBPNet (ATAC-seq), and AlphaGenome (PRO-cap, H3K27ac, H3K27me3), highlighting configuration-specific effects on transcriptional activity and chromatin features. The NFY[-]:8:KLF[+] configuration (second column) represents a novel functional CE predicted by GRACE, whereas the NFY[-]:8:KLF[-] configuration (third column) is a non-functional configuration included to illustrate stereospecificity. **(f)** Example of NFY[-]:8:KLF[+] CE mediated chromatin accessibility across diverse primary cell types, as predicted by AlphaGenome DNase-seq models, demonstrating context-dependent regulatory activity (see **Fig. 5g**).

**Extended Data Fig 6.**
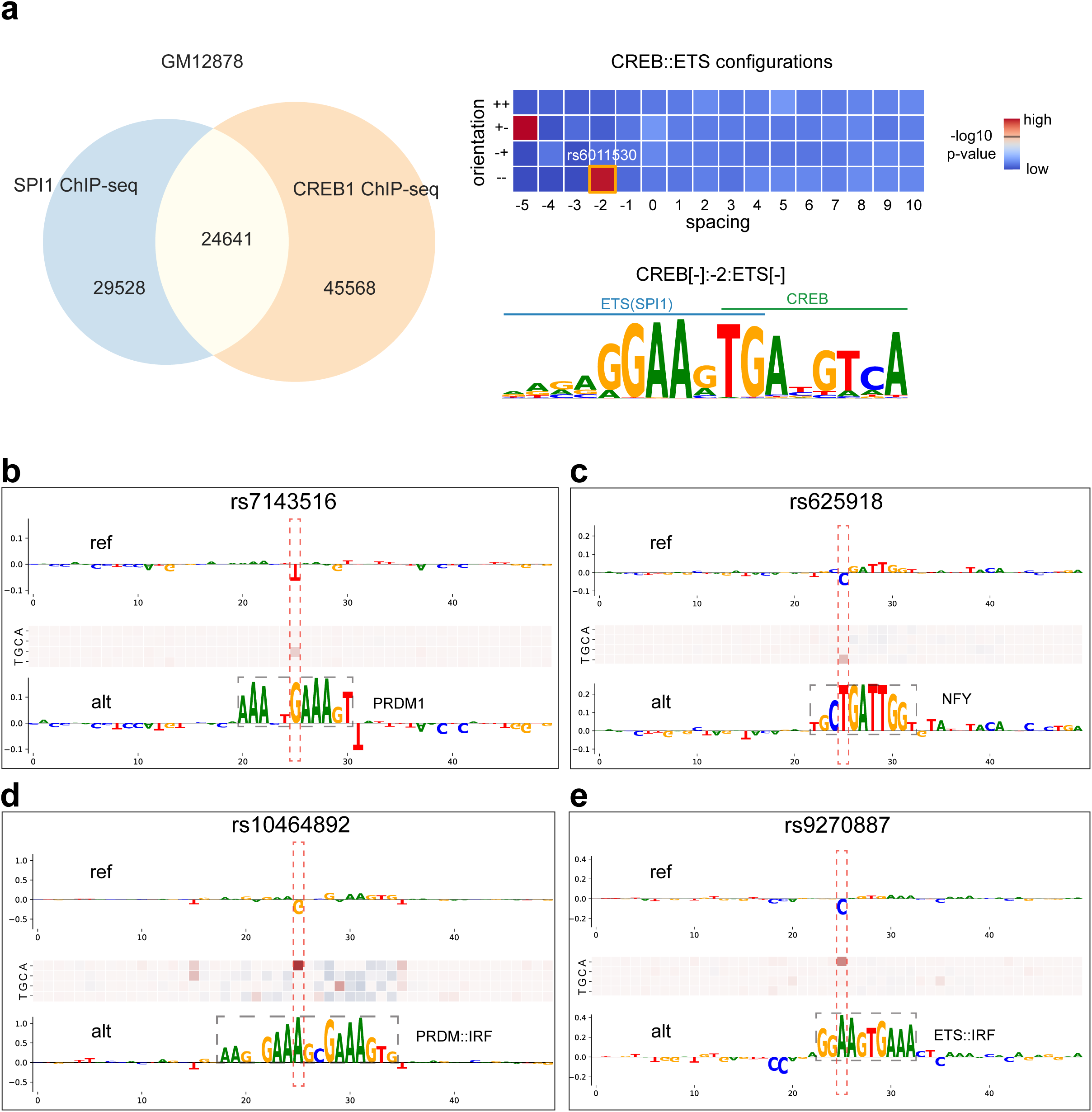
GRACE predictions for additional GWAS variants in autoimmune diseases. **(a)** Identification of a CREB::ETS CE using CEseek and ChIP-seq data for CREB1 and SPI1. The Venn diagram showing the overlap of the CREB1 and SPI1 binding regions (ChIP-seq) in GM12878 cells. Co-binding regions were analyzed with CEseek, using singly bound regions as controls, to identify statistically enriched CREB::ETS configurations (heatmap). One such configuration corresponding to the motif impacted by the GWAS allele *rs6011530* (**Fig. 6e**) is highlighted with an orange box. The corresponding CREB::ETS CE PWM is shown below the heatmap. **(b-e)** All alternative alleles are predicted to result in gain-of-function (GRACE |log2FC| ≥ 0.25). Panels (**b**) and (**c**) show examples of generation of simple TF motifs; panels (**d**) and (**e**) show examples where the alternative allele creates a *de novo* CE configuration. Disease associations are indicated for each variant (see **Supplementary Table 14**). The heatmaps in the middle row depict *in silico* saturation mutagenesis predictions generated by GRACE for the reference sequences.

